# Simultaneous representation of multiple time horizons by entorhinal grid cells and CA1 place cells

**DOI:** 10.1101/2023.04.09.536160

**Authors:** Prannoy Chaudhuri-Vayalambrone, Michael Rule, Marius Bauza, Marino Krstulovic, Pauline Kerekes, Stephen Burton, Timothy O’Leary, Julija Krupic

## Abstract

Grid cells and place cells constitute the basic building blocks of the medial entorhinal-hippocampal spatial cognitive map by representing the spatiotemporal continuum of an animal’s past, present and future locations. However, the spatiotemporal relationship between these different cell types is unclear. Here we co-recorded grid and place cells in freely foraging rats. We show that average time shifts in grid cells tend to be prospective and are proportional to their spatial scale, providing a nearly instantaneous readout of a spectrum of progressively increasing time horizons ranging hundreds of milliseconds. Average time shifts of place cells are generally larger compared to grid cells and also increase with place field sizes. Moreover, time shifts displayed nonlinear modulation by the animal’s trajectories in relation to the local boundaries and locomotion cues. Finally, long and short time shifts occurred at different parts of the theta cycle, which may facilitate their readout. Together, these findings suggest that progressively increasing time horizons of grid and place cells may provide a basis for calculating animal trajectories essential for goal-directed navigation and planning.

## Main

The firing patterns of spatially sensitive cells, such as grid and place cells, are normally viewed as functions of an animal’s current position. However, studies have shown that their firing is often biased towards representing positions the animal has just visited or is about to visit^1–8^. In this way, activity in place cells and grid cells represents the spatiotemporal continuum of an animal’s past, current and future locations. In rats, on average, temporal biases in CA1 place cells tend to be positive and span ∼120 ms^1,6,8^. Previous work suggests that inputs to CA1, representing an animal’s movement in a particular direction, may be used for shifting spatial representations slightly ahead of the animal by defining a small future trajectory segment^5,8–10^. Grid cells, notably, share a constant temporal offset with co-recorded medial entorhinal speed cells^5^. However, the spatiotemporal relationships between different grid modules and between grid cells and place cells have not been established.

Here, we investigated temporal biases in how locations are represented by the medial entorhinal grid cells and CA1 place cells. We found that both grid and place cells predominantly represent space with a prospective bias in a two-dimensional foraging task. Similar to the spatial organisation of the entorhinal grid and hippocampal place cells, which fundamentally depends on the geometry of the enclosure^11–16^, the time shifts of these cell types are modulated by the animal’s trajectory in relation to the enclosure boundaries. In addition, they are also non-linearly modulated by the animal’s locomotion properties, such as speed, angular velocity and acceleration.

Importantly, the magnitudes of these time shifts correlate with the spatial scale of grid cell and the size of the place cell field. Thus, simultaneously recorded grid cells of different grid modules (a grid module is defined as a group of anatomically neighbouring grid cells with similar scale and orientation^17–19^) together with different size place cells may, in principle, provide a nearly simultaneous (occurring within 200 ms; see Methods) readout of multiple time shifts available at any given location. Notably, time shifts in place cells are on average larger than in grid cells. Finally, we found that both grid and place cells with different time shifts tended to fire at different phases of the theta cycle, suggesting that the time horizon of a given cell’s coding window may be organised by the theta rhythm.

## Results

### Place cells have larger time shifts compared to grid cells

To establish how temporal biases are distributed along the dorsoventral axis of the medial entorhinal cortex (mEC) and along the hippocampal-mEC network, we analysed data from a study published by Krupic and colleagues^11^, in which grid cells (n = 249, 6 rats) and place cells (n = 95, 5 rats) were recorded while rats freely foraged in four geometrically distinct enclosures (Fig. 1A). Grid cells and place cells were simultaneously recorded in three of these rats. To investigate whether grid and place cell firing was predominantly prospective or retrospective (or encoding an animal’s current location), we shifted spike times by up to ±2 seconds (in 20 ms steps) relative to the rats’ trajectories. With each time shift, we measured the ‘sharpness’ of the resulting rate map’s firing fields by calculating the spatial variance of its rate map (“zero-lag spatial autocorrelation” ZLAC; see Methods; “lag” here refers to spatial lag). The position of the ZLAC peak nearest 0 ms was taken as the recording’s optimal time shift (Fig. 1B). In this study, positive time shifts indicate prospective firing, while negative shifts indicate retrospective firing.

**Figure 1.**
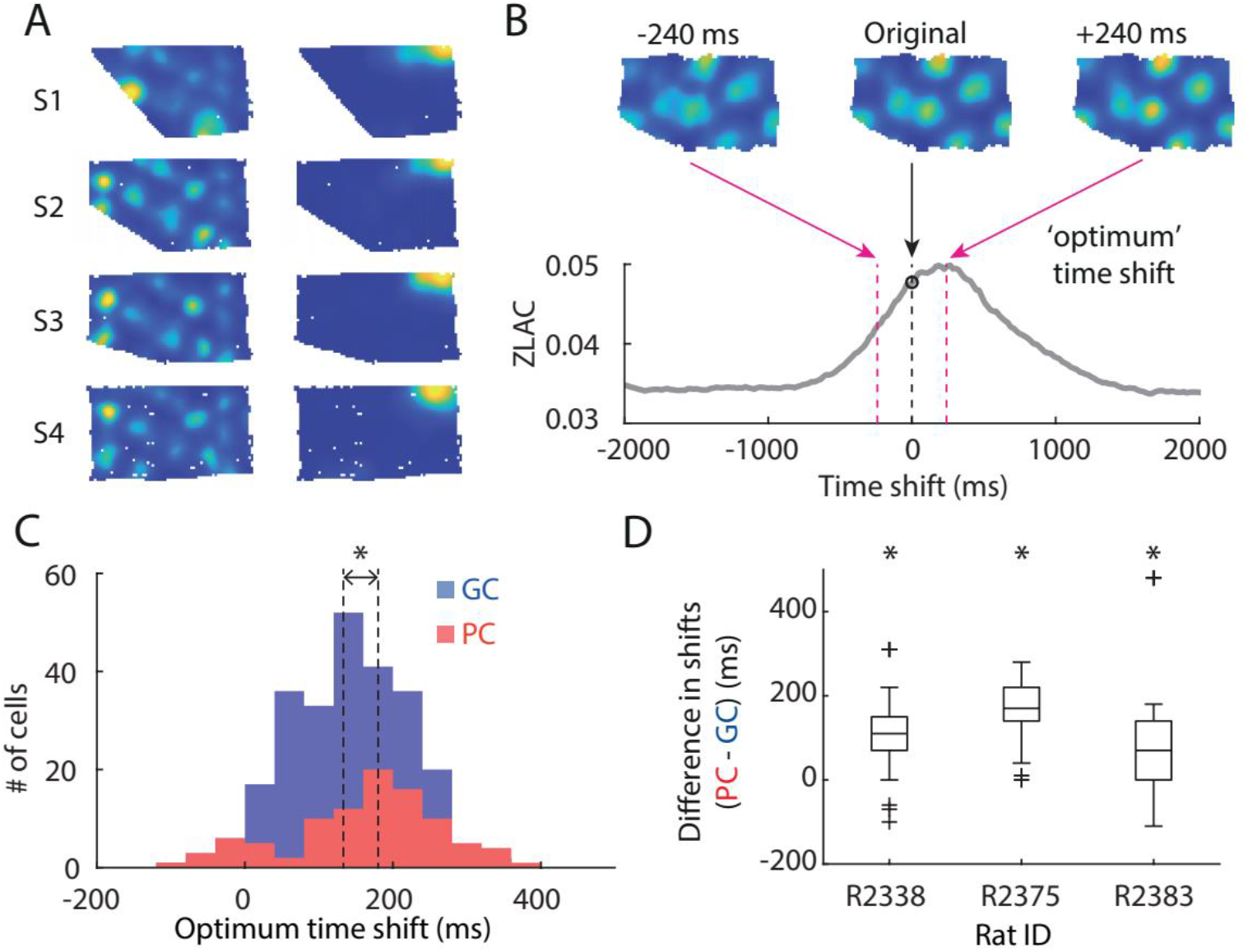
Grid cells and place cells encode space prospectively. (**A**) Rate maps of a typical grid cell (left column) and a place cell (right column) recorded in each of four differently shaped enclosures, “S1”-“S4”. (**B**) Shifting a cell’s spiking location along the trajectory of the rat sharpens firing fields, as measured by the zero-lag autocorrelation. ZLAC: zero-lag autocorrelation. (**C**) Time shift distribution from all 8 rats. This includes each cell’s median optimum time shift across all enclosures. On average, place cells had significantly larger prospective time shifts than grid cells GC: grid cell; PC: place cell. (**D**) The differences between trial-averaged grid and place cell time shifts in each of the three animals with co-recorded grid and place cells (outliers indicated by crosses). On average, recorded place cells showed more prospective time-shift values than grid cells.

Overall, both grid cells and place cells were biased towards prospective firing (Fig. 1C, grid cells (n = 249, blue): median 133 ms (interquartile range (IQR) 116), Z = 13.4, P = 4.5 × 10^-41^; place cells (n = 95, red): 180 ms (IQR 133), Wilcoxon signed-rank test (WSRT): Z = 8.2, P = 2.8 × 10^-16^), with place cells on average having significantly larger prospective time shifts compared to grid cells (Mann–Whitney U test (MWUT): Z = −2.8, P = 4.5 × 10^-3^); N.B. there was no significant difference between the median field sizes of the grid (median 0.22 m², IQR 0.25) and place cells (median 0.22 m², IQR 0.12; MWUT: P = 0.67, Fig. S1). Three rats with co-recorded place cells and grid cells similarly showed significant differences between the average time shifts of the two cell types, with place cells exhibiting larger time horizons (Fig. 1D, MWUTs: 3/3 significant after Bonferroni correction for multiple comparisons, False Discovery Rate (FDR) α = 0.05). In addition to using ZLAC, we calculated the optimal time shifts by maximising spatial information^20^ and peak firing rate, which are commonly used properties for finding optimal time shifts in place cells^4,5,77^. We found that the resultant optimal time shifts were highly correlated (Fig. S2), ruling out the possibility that the results are due to a hidden bias of our novel ZLAC method.

### Enclosure boundaries modulate the optimal time shifts

If time shifts provide a horizon in which the animal’s past, present and future state is being represented, we might expect time shifts to be influenced by boundaries and obstacles that constrain behaviour. Notably, it has been shown that spatial symmetry in grid cells fundamentally depends on the geometry of the enclosure^11,12,15,16^, and grid cell firing resets at the walls^21–23^, which may correct path integration errors and increase animals’ certainty about their current locations. It has been suggested that the amount of information available to the animals may influence their predictive coding horizon: when local cues are abundant, rats show consistent predictive coding, which degrades in cue-deprived environments^1^. We, therefore, hypothesised that the time bias may depend on enclosure shape and the rat’s proximity to the walls.

To address this, we compared the time shifts in four geometrically distinct enclosures, which have been shown to induce local distortions in grid cell firing patterns and local shifts in place fields close to the walls^11^. We reasoned that if time shifts are used to represent immediate future trajectories, which on average, extend further in a larger rectangular (“S4”) enclosure compared to a smaller trapezoidal (“S1”) enclosure, prospective time bias should be larger in the former enclosure compared to the latter.

To address this question, we regressed time shifts against enclosure shape (Fig. 2A; linear regression Δt = *m n* + *b*, where *n* is the enclosure number ranging from 1: most trapezoidal to 4). Consistent with our predictions, grid cell time shifts increased in the larger arenas (inner 95% confidence interval for slope *m*: P_2.5–97.5_ = 3.1–14 ms/arena; cells from all subjects combined). This effect was consistent in direction and magnitude across all subjects (Fig. S3). For place cells, the inner 95% confidence interval for *m* (P_2.5–97.5_ = −16–9.0 ms/arena) included zero, with no consistent effect across subjects (Figs. 2A, S3). Notably, in grid cells, the arena shape remained a minor contributor to the overall time shifts, with the median time shift remaining largely prospective even in the smallest arena (Δt: median 140 ms, IQR 60–200 ms). This implies that arena deformation effects are superimposed on a larger, fixed prospective shift.

**Figure 2.**
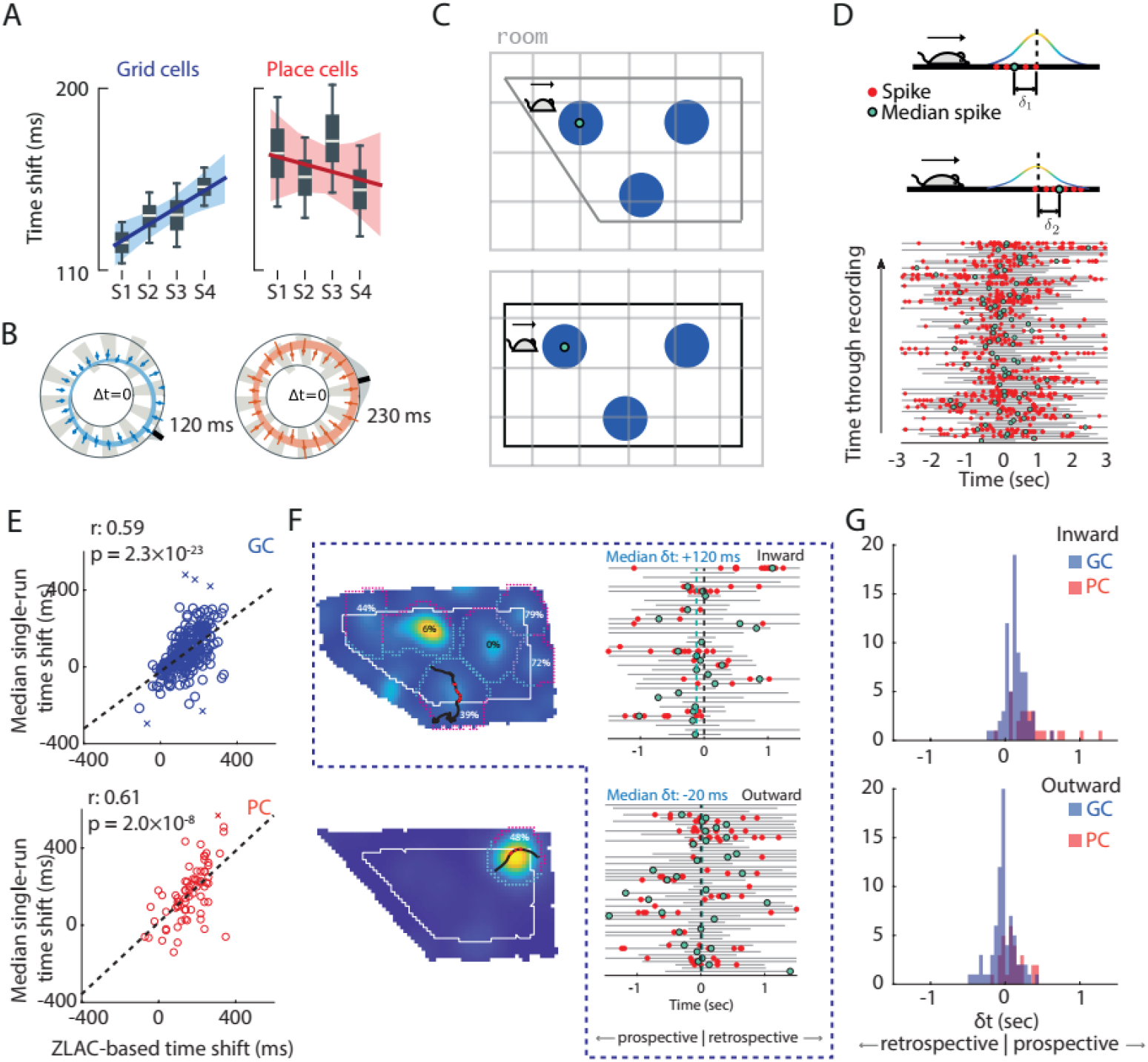
Time shifts depend on external boundaries. (**A**) Changes in time shifts with enclosure shape. Time shifts significantly increased with enclosure size for grid, but not place cells. (**B**) Time shifts are consistently larger for eastward headings. Log-polar plots show heading on the angular axis and Δ*t* on the radial axis (Inner axis ring: Δ*t* = 0; Coloured rings: 95% confidence intervals for polar least-squares regression; Points: median Δ*t* in 15° bins with (bars) 95% confidence). The exterior black bar indicates the regressed direction of largest Δ*t* (“*θ*_max_”). Grey-shaded regions on the exterior reflect a von Mises model of uncertainty in *θ*_max_. All plots reflect single-run time shifts aggregated over cells from all subjects after removing per-cell variability. (**C**) Schematics of a rat heading eastwards. Grid fields (blue) close to the west wall shift westwards in the rectangle compared to the trapezoid. These shifts are consistent with time shifts becoming more prospective when the rat is moving eastwards. Grey: room frame of reference. The shifts are shown not scale to facilitate visualisation. (**D**) Schematics (left) and a plot (right) showing the timing of individual spikes (red dots) and median spikes (green dots) on all ‘valid’ runs for a particular grid cell recording. Note that several median spikes represent the only spike fired on a given run. (**E**) Scatter plot comparing ZLAC-estimated fixed time shifts with median single-field-run time shifts. Each point represents the mean measure for each cell (249 grid cells, 95 place cells). There is a significant positive correlation between the two measures. Crosses (×) mark outliers calculated from cells without enough valid runs. (**F**) Left: Each rate map was split into inner and outer zones of equal area (white line, solid) based on the relative occupation in each zone (“peripheral” if ≥30% of its area fell in the outer zone). Runs were deemed “inward” if they started on the outer half and ended on the inner half of each field and “outward” if vice versa. Black trajectory: an inward run; red dots indicate spikes fired along this run. Right: inward and outward runs from the grid cell rate map shown, and the spikes fired on each run. Dashed green and black lines indicate grand median single-run time shift and zero time shift: positive on the top and slightly negative in the bottom. (**G**) Median spike times on inward and outward runs across all 73 grid cells with sufficient runs.

Next, we asked whether there was a bias in time shifts depending on the animal’s running direction. We used the animal’s movement direction (‘heading’) as a correlate of the animal’s past trajectory^22^ (i.e. the animal’s going eastwards approximates a recent encounter with the west wall, whereas the animal going southwards suggests its recent encounter with the north wall etc.; N.B. heading and head direction are highly correlated and show consistent results; Fig. S4). We used a polar regression Δ*t* = *d* cos(*θ* – *θ*_max_) + *c*, where *d* captures the depth of directional modulation, and *θ*_max_ captures the direction in which time shifts are most prospective. For both grid cells and place cells, time shifts showed significant prospective heading modulation (modulation depth spanning *d* = 46–87 ms in all subjects for grid cells and *d* = 62–280 ms in 3/5 subjects for place cells, FDR TSBH α = 0.05), with the largest time horizons consistently reflecting eastward directions (Figs. 2B, S5 *θ*_max_ ranges over all subjects: heading GC: S82°E–S30°E, PC: N56°E–N77°E). Further investigation revealed that directional modulation depended on the arena shape, showing a preference for eastward directions in the rectangular arena and south-eastward directions in the trapezoidal arena. Such directional bias suggests that the animals may be using the frame of reference associated with the external room instead of the internal walls of the enclosure to estimate the field locations. Namely, the rat reaches fields sooner than expected when it travels from west to east in the rectangular enclosure compared to the trapezoidal enclosures. On the other hand, in the trapezoidal enclosure, the fields, on average, may be ‘squeezed upwards’ as a result of deformations and the animals will tend to show more south-east directed bias in time shifts (Fig. 2C).

Finally, we asked whether the time shifts depended on the animal’s trajectories in relation to the enclosure walls. We hypothesised that time shifts in grid cells and place cells may be modulated by proximity to the walls, which act as reset points for updating the grid cell firing^21^–^23^ to reduce the accumulation of path integration errors (hence decreasing ambiguity about the location, which is known to affect time shifts in place cells^1^). Furthermore, such modulation should change depending on whether the animal approaches or moves away from the wall. We reasoned that if time shifts indeed reflect the look-ahead predictive time horizon of the medial entorhinal-hippocampal network, they should be shorter when the animal approaches the wall compared to when it is leaving the wall. To investigate this, we first divided each grid and place cell rate map into individual firing fields and used these to segment the animal’s trajectory into individual runs through single fields^24^. For each spike train associated with a single run, we calculated a ‘single-run’ time shift “δ” as the time difference between the “median spike” (Fig. 2D, green dots) and the point on the run closest to the field centre (positive shifts = prospective coding; N.B. often the median spikes can be the only spikes). We confirmed that using this alternative method, each cell’s grand median single-run time shift (i.e. the median of each recording’s median single-run time shift) was significantly correlated with the cell’s ZLAC-optimal time shift (Fig. 2E, grid cells (n = 242 excluding outliers): Pearson’s r = 0.59, P = 2.3 × 10^-23^; place cells (n = 92, excluding outliers): Pearson’s r = 0.61, P = 2.0 × 10^-8^).

We then compared these single-run time shifts on runs towards and away from the walls (“outward” and “inward” respectively) in fields near the walls (Fig. 2F-G; see Methods). On inward runs (away from the walls), grid cells showed prospective time shifts (Fig. 2G, n = 73 cells with sufficient runs: median 132 ms (IQR 167); WSRT: Z = 6.3, P = 2.8 × 10^-10^), as expected based on grid cells’ overall average time shifts. However, outward runs (i.e. when the animal approached the wall) had significantly shorter time horizons (difference between time shifts of inward and outward runs in each map: median 130 ms (IQR 214), WSRT: Z = 5.7, P = 1.4 × 10^-8^); and were neither prospective nor retrospective (median 25 ms (IQR 156); WSRT: Z = −1.6, P = 0.10). Place cells showed similar results: inward runs tended to have larger prospective shifts than outward runs (difference between time shifts of inward and outward runs in each map: median 260 ms (IQR 281), n = 29 place cells with sufficient runs, WSRT: Z = 3.7, P = 2.3 × 10^-5^). However, for place cells, both inward and outward runs showed significant prospective time shifts (inward runs: median 300 ms (IQR 310), WSRT: Z = 4.6, P = 4.8 × 10^-6^; outward runs: median 70 ms (IQR 184), WSRT: Z = 3.0, P = 2.6 × 10^-3^).

### Nonlinear modulation of time shifts by internal cues

To dissociate the influence of the wall from behaviour correlated with turning or stopping, we checked for systematic differences in behaviour between inward and outward runs near walls. Consistent with rats slowing as they approach the wall, acceleration was more positive in inward runs for both cell types (median difference: GC: 0.035 m·s^-2^, IQR 0.054, n = 73 cells with > 15 runs; PC: median difference 0.044 m/s^-2^ IQR 0.044, n = 29 cells with > 5 runs), significant in both cases (GC: p = 3.9 × 10^-13^, PC: p = 2.6 × 10^-6^; WSRT). The median angular velocity was similar between inward and outward runs in the data for grid cells (WSRT: p = 0.91), but significantly different for place cells (median difference 4.3 °/s (IQR 18.2°), WSRT: p = 7.7 × 10^-4^). Given these behavioural biases, we tested whether time shifts depended on behaviour more generally. We measured the rank correlation between per-run time shifts and acceleration, forward speed, and angular speed (Methods). We converted data to ranks on a per-recording basis before combining to remove per-cell and per-rat effects.

Acceleration was significantly (after α = 0.05 TSBH FDR control) positively correlated with single-run time shifts for both cell types in all subjects save one (n.s for place cells in R2377), with correlations spanning ρ = 0.08 – 0.12 (Figs. 3A, S6); the plots show the result of the rank regression after converting back to physical units by inverting the rank transform (Methods). Positive accelerations correspond with more prospective shifts and negative accelerations to smaller (and sometimes negative) time shifts. This effect saturated for the most extreme positive or negative accelerations and showed an inflection point around zero. This suggests an almost binary switch between more prospective coding when accelerating and less prospective coding when decelerating.

**Figure 3.**
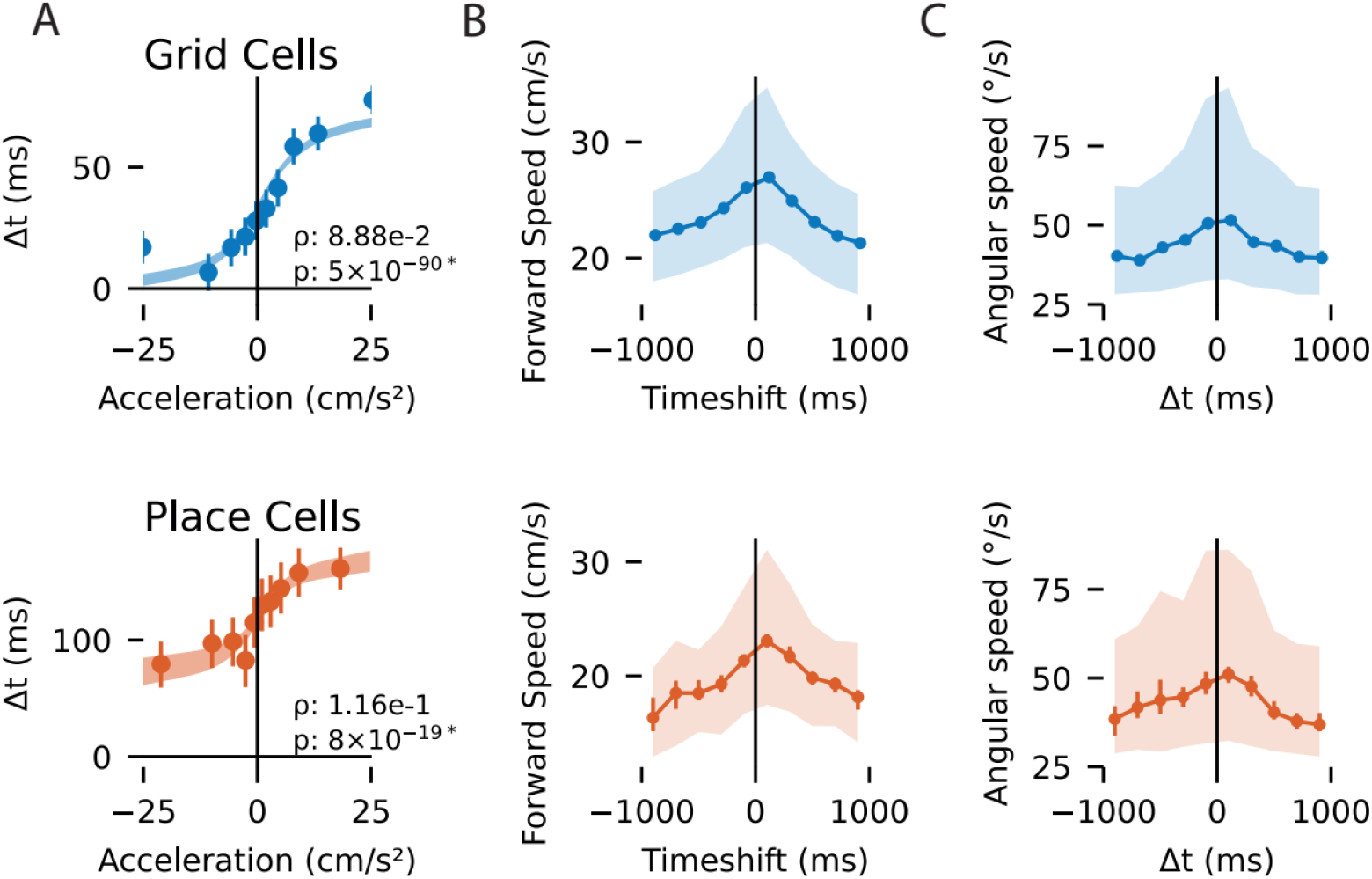
Nonlinear relationships between time shifts and behaviour for place (red, top) and grid cells (blue, bottom). (**A**) Positive accelerations correlate with more prospective time shifts (Δt). Plots show the estimated linear rank regression of time shifts given acceleration after inverting the rank transform back to physical units (shaded: 95% confidence of regressed fit; points: mean Δ*t* within each decile; lines: 95% confidence). (**B**) The largest positive and negative time-shifts are associated with slower running speeds (consistent across subjects, see Fig. S7; Points: median within histogram bins; Bars: 95% confidence in the median; Shaded: inter-quartile range within each bin). Faster running speeds are associated with more typical time shifts (c.f. Fig 1C). (**C**) The relationship between time shifts and the angular speed of head directions was similar to that of forward speed, albeit noisier.

Consistent with previous works^3–5^, correlations between time shift and either forward or angular speed were small (forward speed, GC: ρ = −0.06–0.01, PC: ρ = −0.05–-0.00; angular speed, GC: ρ = −0.07–0.02, PC: ρ = −0.07–-0.02) and not significant in individual rats. However, these correlations were negative in a majority of subjects and reached significance when aggregating all runs across rats (forward speed: GC: ρ = −0.006, PC: ρ = −0.03; angular speed: GC: ρ = −0.02, PC: ρ = −0.04). A closer inspection revealed a nonlinear relationship (Figs. 3B-C, S7), whereby the average forward speed (and angular speed) was consistently higher on runs with time shifts closer to zero and runs with either very large positive or negative time shifts were associated with slower average forward and angular speeds. This finding is consistent with the idea that time shifts may reflect the information about the future and past actions usually occurring when the animal is relatively immobile^25,26^.

To check whether these behavioural correlations with time shifts might be related to boundary effects or arena shape, we compared rank correlations between three pairs of topographic subsets of the data: arenas “S1” (trapezoid) vs “S4' (rectangle), near walls vs interior regions, and the right vs left halves of the enclosure. For acceleration and speed, correlation coefficients never differed more than expected by chance for more than one subject for any comparison (z-test; TSBH FDR α = 0.05). A more detailed comparison found no consistent topographic dependence for time-shift–acceleration correlations (Fig. S8), and differences in directional modulation *d* were never significant for any comparison (both heading and head direction; two-tailed test on the bootstrapped difference in *d*; TSBH FDR α = 0.05). No significant regional differences in the correlations with acceleration or speed emerged when aggregating cells from all rats.

Finally, we checked whether the tendency toward larger time shifts in the larger arenas could be attributed wholly to behavioural correlations. We measured the partial correlation between arena shape (S1–S4) and time shift (ranks), conditioned on acceleration (ranks) and heading angle (cos(*θ*), sin(*θ*)). This summarises the linear variation in (ranked) time shifts that can be uniquely explained by, e.g. arena shape and no other variables (N.B. for simplicity, we omitted forward and angular speed analysis due to their substantially lower nonlinear influence on time shifts, see above). The positive trend in grid cells persisted (ρ_partial_=0.029, p<10^-11^; c.f. raw correlation ρ_0_=0.030; single-field runs from all subjects aggregated). Likewise, significant partial correlations between run direction (near walls) and time-shifts persisted (GC: ρ_partial_=0.105, p<10^-27^, c.f. ρ_0_=0.145; PC: ρ_partial_=0.148, p<10^-6^, c.f. ρ_0_=.221; single-field runs from all subjects aggregated). Finally, significant (albeit smaller) partial rank-correlations persisted for (acceleration and time shift) when conditioning on run-direction near walls (GC: ρ_partial_=0.068, p<10^-11^, c.f. ρ_0_=0.109; PC: ρ_partial_=0.095, p_<_10^-3^, c.f. ρ_0_=0.175; single-field runs from all subjects aggregated).

In summary, time shifts are generally more prospective in the larger arenas when running inward from the wall, when accelerating (vs decelerating), and when moving eastwards (vs westwards). Acceleration and inward/outward run direction are correlated but not wholly redundant. Heading and arena-shape correlations potentially reflect the effects of room distortion, as previously explored in^11,22^.

### Time shifts are correlated with the spatial scale of firing patterns

Grid cells are topographically arranged into distinct grid modules along the dorsoventral axis of the mEC, with smaller scales located dorsally and larger scales at ventral parts of the mEC^17^–^19,27^ and were shown to exhibit independent neural processing^18,28^. Hence, we asked whether place and grid cells’ time shifts also varied with their spatial scale (Fig. 4). We reasoned that if larger grids represent larger distances, they may also exhibit longer time horizons. For this analysis alone, we added a separate dataset recorded in larger enclosures (∼2.8 m × 3 m, see Methods), which included grid cells with much larger scales. We found a significant positive correlation between grid scales and time shifts, with larger-scale grid cells having more prospective time shifts (Fig. 4A-B, n = 249 + 21 cells from small + large enclosures respectively: Spearman’s ρ = 0.44, P = 5.3 × 10^-14^). Similarly, place cell optimum time shifts were positively correlated with their field sizes (Fig. 4C-D, n = 74 + 24 cells from small + large enclosures, respectively: Spearman’s ρ = 0.54, P = 1.3 × 10^-8^).

**Figure 4.**
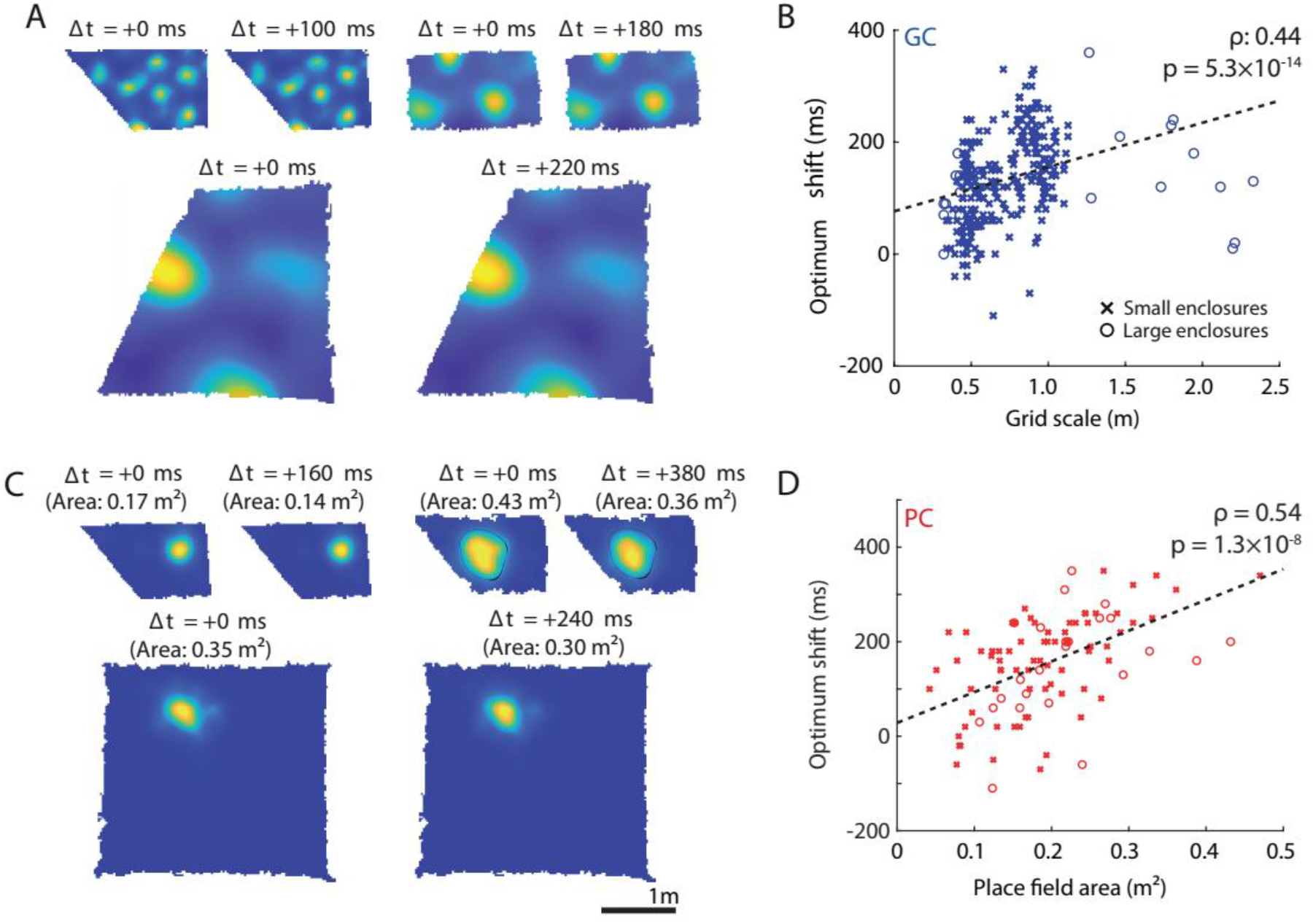
Time shifts in grid and place cells are correlated with firing field size. (**A**) Three example grid cells with different scales (upper left: 0.45m, upper right: 0.92 m, lower: 2.10m, in the large trapezoidal enclosure described in Methods), before and after time shifting (Δt: applied time shift). Note that these examples, while representative of optimum time shifts at those scales, result in very small visual changes to each firing rate map. (**B**) There is a significant positive correlation between grid scale and optimum time shift. (**C**) Three example place cells with different firing field sizes, before (left) and after time shifting (right). Top right field is outlined with a blank line to facilitate visualisation of the decrease in the area with time shifts. (**D**) There is a significant positive correlation between place cell firing area and optimum time shift. Linear regression was used to draw a line of best fit (dashed black). The same scale bar is used for all the rate maps.

To exclude the trivial explanation that these correlations resulted from a methodological bias towards estimating larger time shifts in larger fields, we repeatedly simulated grid cells and place cells of different scale and field sizes, respectively, with zero underlying time shifts, matching the recorded data sets (Fig. S9, see Methods). In simulated maps with larger grid scales/place field sizes, we found that the optimum time shift estimates had a higher variance, but the means were not significantly positive or negative. This suggests that our methods do not introduce bias towards identifying more positive time shifts in larger-scale firing rate maps.

### Grid modules provide multiple simultaneous time shifts

We next asked whether on a moment-by-moment basis overlapping fields of co-recorded grid cells from the same grid module had similar time shifts, whereas cells from different co-recorded grid modules consistently showed larger time shifts. This would suggest that the time shift is signalled at the grid module level and that a spectrum of multiple time shifts may be read out nearly simultaneously (within < 200 ms time window; see Methods) from the outputs of different scale grid cells. To address this, we again segmented the animal’s trajectory into individual runs through single grid fields^22^ and calculated the difference in “δ” time shifts between overlapping individual runs for all simultaneously recorded grid cell pairs. Indeed, we found that the difference between the optimal time shifts was not significantly different from zero in co-recorded grid cells of the same grid module (Fig. 5A, mean ± s.e.m.: −2.3 ± 5.0 ms, t_6249_ = −0.46, P = 0.64, Student’s t-test). However, such difference in time shifts was consistently positive in larger grid modules (Fig. 5B, mean ± s.e.m.: 78.8 ± 6.9 ms, t_10101_ = 9.70, P = 3.64 × 10^-22^, two-sample t-test) indicating that multiple progressively increasing time horizons within a few hundreds of milliseconds range can be readout nearly simultaneously at any given point in the environment (N.B. the co-recorded grid fields of both larger and smaller scales as well as place fields were distributed across the entire enclosure).

**Figure 5.**
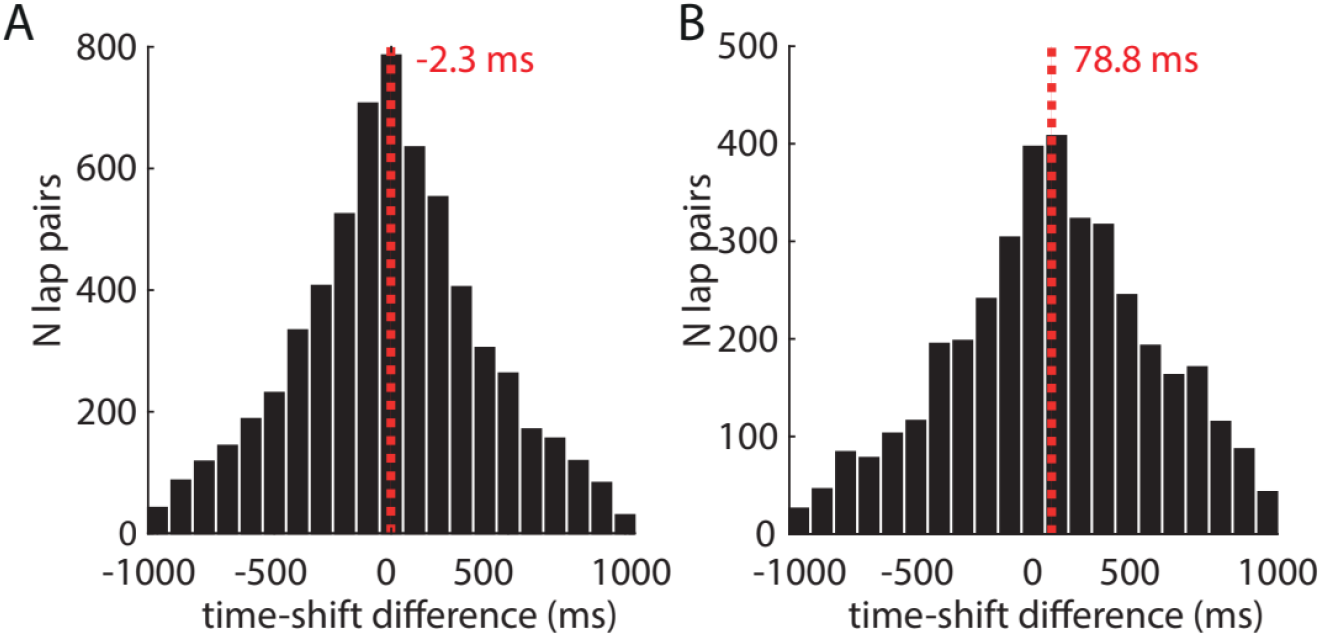
Distinct grid modules provide a nearly simultaneous readout of gradually increasing time shifts. **(A)** Histogram showing the distributions of time shift differences between all grid cells from the same modules. (**B)** Histogram showing the distributions of time shift differences between all grid cells from different modules. Dashed red lines show average time shift differences.

### Time shifts show the differential distribution across theta phase

Cell activity across the hippocampal formation is organised in relation to the theta rhythm^29–33^. Place cells in CA1 tend to fire near the trough (phase 0°) of the LFP theta oscillation in *stratum pyramidale*^29,30,32^ and continue spiking at progressively earlier phases of each theta cycle as the animal traverses a cell’s firing field (a phenomenon known as phase precession^20,31^). Both grid and place cells can show theta precession^20,31,34^. In grid cells, theta modulation is layer dependent^34^ with mEC layer II grid cells often phase precessing, whereas layer III grid cells showing both theta precession as well as theta locking activity (i.e. the tendency to fire at a specific theta phase). Furthermore, experimental evidence shows that several variables correlate with the LFP theta phase, including the relative inputs that CA1 receives from the medial entorhinal cortex and CA3^29,32,35^; the ease of inducing long-term potentiation/depression at CA3 synapses to CA1^36,37^; and the timing of fast vs. slow gamma oscillations in CA1^29^. Such observations support models in which the encoding and retrieval of spatial memories (e.g., associations between specific locations and food reward) occur on separate phases of LFP theta^38^. In such a context, it is possible that each cell’s prospective bias during a specific trial is correlated with the mean theta phase at which it tended to fire, reflecting the balance between that cell’s involvement in encoding and retrieval during the trial.

To explore this, we looked at place and grid cell firing with respect to the theta rhythm in CA1. Previous studies have shown that the trough (phase 0°) of the theta rhythm in stratum pyramidale of CA1 aligns with the phase associated with the highest probability of CA1 place cell firing^29,30,32^. Hence, for each rat, we shifted the measured theta phase such that 0° corresponded to this peak in CA1 activity (Fig. S10). This accounted for differences in recording location within each rat and enabled closer comparison with previous studies. To allow a comparison of place cells with grid cells, we also analysed grid-cell firing in terms of its relationship to the stratum pyramidale theta rhythm rather than the entorhinal theta rhythm. After alignment, we found that CA1 place cells spiked maximally at 0.0 ± 72.0° (circular mean ± circular standard deviation), while grid cells spiked maximally at 268.4 ± 78.1°, with place-cell theta phases lagging grid-cell theta phases (Fig. 6A-B), consistent with cell firing probabilities of principal cells in CA1 and mEC layer II^31^.

**Figure 6.**
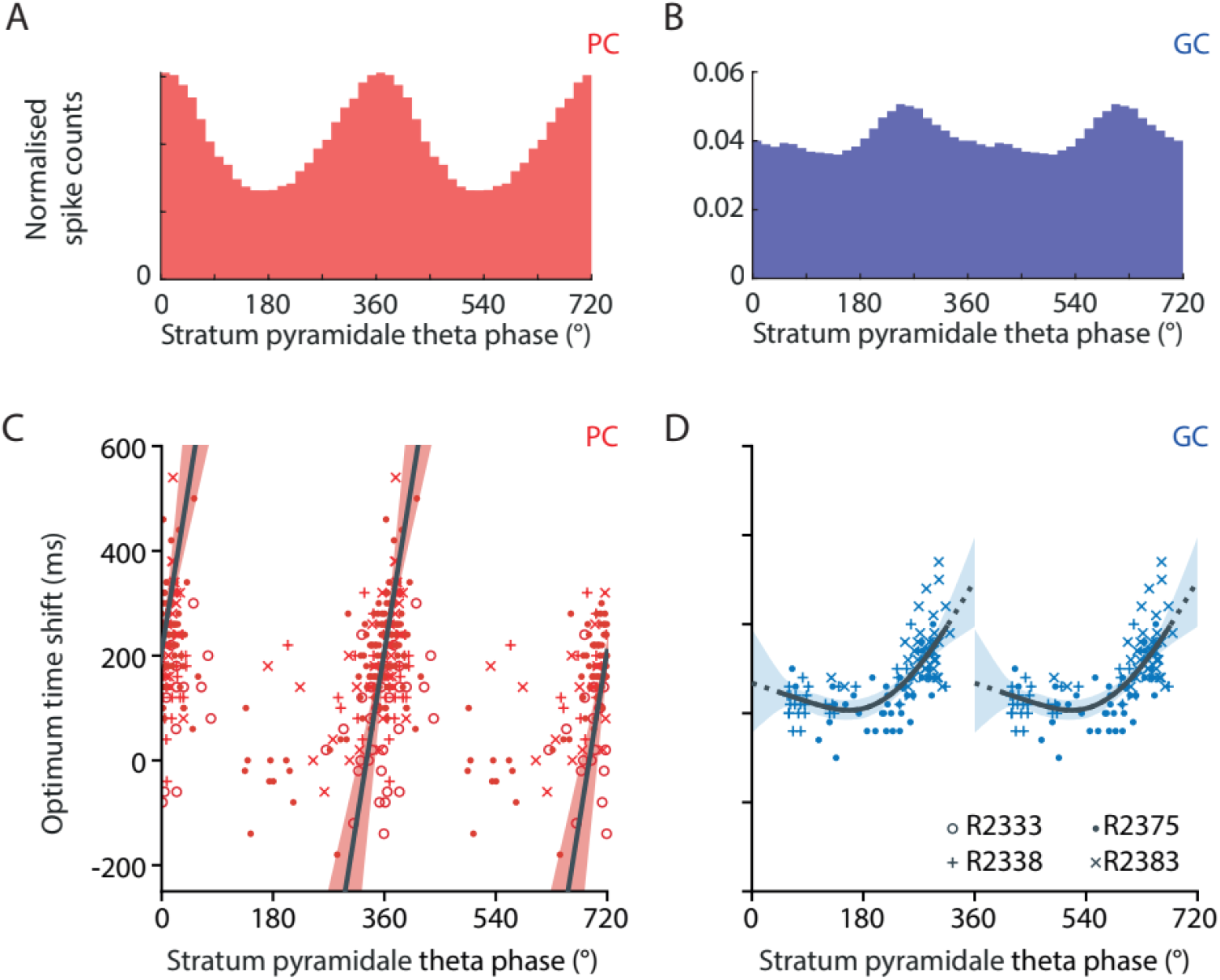
Time shifts in place and grid cells are correlated with their preferred CA1 pyramidal theta phase. Histogram showing normalised spike counts of place cells (**A**) and grid cells (**B**) with respect to stratum pyramidale theta phase. Scatter plot showing each place cell (**C**) and grid cell (**D**) recording’s preferred theta phase against its optimum time shift. PC: place cell; GC: grid cell Grid cells showed a nonlinear relationship saturating at lower preferred theta phases. Trend lines (black) reflect a circular–linear regression for place cells (C) and a cubic-spline regression for grid cells (D; four basis functions spaced equally over 0–360°). Shaded regions show the 2.5–97.5th percentile confidence intervals (bootstrap; 1000 samples). Only cells showing significant theta-phase coupling are included.

We next asked whether each cell’s preferred firing phase relative to the theta rhythm was correlated with the size of its prospective time shift. Several place cells (28/56) and grid cells (29/39) showed significant theta precession (Methods). We could not detect a significant difference between the time shifts of precessing and non-precessing cells (Table S1; MWUT: place cells: P = 0.23; grid cells: P = 0.31). Hence, to investigate the relationship between average time shift and theta phase at a single cell level, we limited this analysis to cells with statistically significant theta phase locking, regardless of whether they showed phase precession (Methods). We found that both place and grid cells showed a significant positive correlation between their average preferred theta phases and their optimum time shifts (Fig. 6C; place cells, n = 239 recordings: Kempter circular-linear correlation coefficient (CLCC) = 0.36, P = 3.6 × 10^-6^; Fig. 6D: grid cells, n = 117 recordings: CLCC = 0.65, P = 3.3 × 10^-11^). This was also true at the level of cells from individual rats (Table S2). Curiously, grid-cell time shifts showed a nonlinear relation with the theta phase, saturating at small values for cells with earlier preferred theta phases (Fig. 6D). To summarise this trend, we regressed a piecewise linear model Δt = max[Δ_0_, *m* · (*θ* − *θ*_0_)] to the combined grid-cell data from all animals. Bootstrap confidence intervals suggest that grid-cell time shifts saturate around Δ_0_ = 19 ms (median; IQR 8.8 ms), with the linear trend beginning around *θ*_0_ = 220° (median; IQR 11°). Overall, these findings suggest that the horizon of a cell’s prospective coding may depend on the phase of its coupling to the hippocampal theta rhythm in agreement with previous studies^2,29,39^. This could facilitate the readout of multiple increasing time horizons from grid cells of different scales (Fig. S11) and place cells of different place field sizes.

It is important to note that this suggested correlation is not a necessary consequence of theta precession. In theta precession, cells spike at progressively earlier phases of each theta cycle as the animal traverses a cell’s firing field. At the level of an individual cell, therefore, precession does predict that spiking at later theta phases is technically more “prospective”. However, precession only describes this gradient in phases for an individual cell^20,40^. It does not constrain the mean phase at which each cell fires. Hence, different cells can display firing locked to different mean phases (though these mean phases are usually clustered around 0°), regardless of whether they precess.

## Discussion

Here we show that time horizons (i.e. the average look-ahead time range) of the grid and place cell positively correlate with their scale and field size, respectively, enabling nearly simultaneous readout of a spectrum of progressively increasing time horizons over a few hundred milliseconds. Distinct grid modules, whose scales increase along the dorsoventral mEC axis^17–19,27^, may provide a nearly instantaneous readout of future positions at multiple time horizons to hippocampal place cells. In line with this hypothesis, our findings suggest that place cells tend to be more prospective than their field-size-matched grid cells, potentially reflecting the contribution of inputs from larger grid cell modules^41–43^. Furthermore, in our dataset, the firing of CA1 place cells appears to lag that of mEC grid cells in the theta cycle (Fig. 6), which may reflect the flow of information from grid cells (likely mostly recorded in mEC layer II/III based on post-hoc histology^11^) to CA1 place cells.

Previously it has been suggested that observed prospective time shifts may reflect the animal's 'true' location, positioned under its nose^1^. In contrast, our data show that place and grid cells simultaneously encode a continuum of locations, which range from the centre of a rat's head to slightly more forward from its nose (see Fig. S12 for median speed distribution) and are determined by the scale and the field size of grid and place cells, respectively, as well as modulated by multiple external and internal cues such as boundary conditions, heading direction, acceleration, forward and angular speeds. The range of simultaneously encoded locations hints that these time shifts may not simply represent the rat’s ‘true’ single location, but instead, they may reflect ongoing computations by the entorhinal-hippocampal network, which take into account past, present and future states. We found that similar to spatial grid structure, the time horizon fundamentally depends on the external boundaries of the enclosure^11,12,15^. In the most polarised trapezoid, the time horizon shrinks compared to a rectangular enclosure suggesting that the time horizon expands when animals expect a longer run path. This is further corroborated by time shifts showing a significant positive bias in a direction towards the unchanging part of the environment (eastward). Furthermore, information from the boundaries may reduce the overall uncertainty of the rat’s whereabouts, resulting in shorter time horizons as the rat approaches the boundaries. In addition to the effects of external cues, internal cues also play a role in modulating the time horizons with a nonlinear relationship. Namely, time shifts show sigmoidal dependence on acceleration. Previously, positive acceleration was associated with an increase in theta frequency^44^. Theta frequency is associated with the pacemaker of entorhinal-hippocampal computations^2,20,29,32,33,38,40,45,46^, thus this increase would suggest that larger time horizons (i.e. further look-ahead mode) may be used in more rapid entorhinal-hippocampal computations during accelerated movements and may provide a more efficient way for error correction due to rapid acceleration. At the same time, larger time shifts at lower angular and forward speeds are consistent with the idea that time shifts may reflect the information about the future and past actions usually occurring when the animal is relatively immobile^25,26,4747^. It is important to note that our data suggests that these internal and external factors are not entirely redundant and may be shaping time shifts independently of each other. Moreover, the combined influence of all these factors cannot fully explain the observed positive time shift bias, suggesting that it may be an intrinsic property of the entorhinal-hippocampal network.

Our second major finding provides compelling evidence that hippocampal theta oscillations may coordinate the readout of the spectrum of locations encoded by multiple place and grid cells. We found that spiking at later stratum pyramidale theta phases is associated with more prospective time shifts in both grid cells and CA1 place cells. While in place cells, this association appears to be linear, in grid cells, it is strongly nonlinear, hinting towards the possibility that distinct grid modules may be providing inputs at different theta phases (Fig. S11). Furthermore, it has been previously suggested that information processing by the entorhinal-hippocampal network is organised with respect to theta and gamma oscillations^2,20,29,32,33,38,40,45,46^. Namely, the prospective coding in CA1 place cells tends to be accompanied by slow gamma oscillations, whereas fast gamma oscillations are associated with more retrospective coding^2^. Fast and slow gamma were shown to occur at different theta phases associated with distinct modes of communications between the CA1-mEC and CA1-CA3^29^, which may correspond to memory encoding and retrieval modes respectively^38^. Hasselmo&Eichenbaum^48,49^ explicitly predicted a nearly simultaneous representation of sequential activity in the hippocampus and the superficial layers of the medial entorhinal cortex, with the timing of such activity based on the hippocampal theta phase. Moreover, subsequent computational models predicted that such sequences should include place cells with multiple place field sizes in order to support ‘flexible’ navigation to the goals^47^ (including distal goals). Our findings provide some compelling evidence supporting such models and further extend them. Namely, the multi-size-field sequences were primarily modelled for place cells. We show that this applies to grid cells as well. We argue that grid cells may represent a more robust system than place cells since they maintain their ‘canonical’ scales invariable across different experimental conditions^27^ and, therefore, may be read out using a consistent relationship to theta phase. Moreover, we found that grid cells may encode the time shifts on a module basis, which may help to decrease the accumulation of noise. Specifically, each module has many colocalised ‘identical’ grid cells, whose sum input would be less noisy, while multiple grid scales of each module would ensure that the distribution of these look-ahead representations is sufficiently dense. Both of these features were identified as crucial for reliable navigation to distal goals^47^.

Finally, it is important to stress that in these computational models, ‘decision points’ during navigation to a goal spanned much wider look-ahead ranges^47,48,50,51^. Place cells’ replay activity^52–54^, which occurs when the animal is relatively immobile, was suggested to correspond to such look-ahead sequences^47,51^. Here, we show another type of sequential firing in place cells and grid cells, which is continuously present, including when the rat is moving and spans much shorter ranges, which depend on the rat’s acceleration, angular and forward speeds as well as the rat’s relations to the boundaries and the shape of the enclosure. It is possible that introducing a goal would modulate time horizons in line with observations that goals may significantly influence place^55,56^ and grid^57,58^ fields. The successor representation hypothesis^59^ predicts that one role of grid cells may be to transform distal goals into immediate movement commands. This requires information about both the position and the desired changes in position (position derivative). By dynamically combining inputs from different grid modules with progressively longer time horizons, hippocampal place cells may be able to generate smooth future trajectories ahead of the animal, which may provide the basis for goal-directed navigation and planning^56,59^.

## Methods

This work was conducted in accordance with the UK Animals (Scientific Procedures) Act (1986). The following sections summarise the acquisition of the original data from^11^ and previously unreported data from 7 rats recorded in larger enclosures. Below we describe the analytical methods applied to the data in this study.

### Animals

10 adult male Lister Hooded rats were chronically implanted in the left and/or right hemisphere with a microdrive (Axona) loaded with four tetrodes (9 rats), or a single Neuropixels probe^60^ (1 rat; R2405). Tetrodes were aimed at the superficial layers of the medial entorhinal cortex (mEC, 6 rats: 4.3–4.5 mm lateral to the midline; 0.2–0.5 mm anterior to the sinus; angled forwards in the sagittal plane at 0–10° and 1.5 mm below the pia) and/or CA1 region (6 rats: 2.5 mm lateral to the midline; 4 mm posterior to bregma and 1.4–1.8 mm below the pia). See Supplementary Information Fig. S1 in^12^ for more details and histology results. In short, in most of the cases the recordings were done from layer II or layer III of mEC.

Neural activity was recorded while the rats foraged for food in four familiar polygonal enclosures (Fig. 1A), which varied in shape from a rectangle (poly180°, S4; 1.8 m × 1.0 m) to a left trapezoid (poly129°, S1) and were presented in random order.

The rat’s position was determined based on an array of two infrared light-emitting diodes (LEDs) fixed to the rat’s head and recorded with an overhead infrared camera (sampled at 50 Hz). The LEDs were fixed to the plastic screw post cemented to the head of the animal ∼3 mm posterior from the bregma and ∼3 mm lateral (always on the left-hand side) from the midline. The position of the plastic screw post corresponds to the position of the rat at zero time shift.

To analyse the relationship between firing scale and time shifts over a larger range of grid cell scales (Fig. 4), we included electrophysiological data collected from 7 rats during a separate series of experiments in larger enclosures. During these experiments, rats foraged in two polygonal enclosures, which differed only in the configuration of their western wall: a 2.8 m × 3 m (north-south × east-west) rectangular enclosure (Fig. 4C) and a right trapezium (Fig. 4A; the northwestern corner is displaced by 1.2 m horizontally between the two enclosures). The rats were trained for 3–4 weeks to forage for sweetened rice randomly scattered throughout these enclosures. Subjects were then implanted with a pair of 8 tetrodes (2 rats) or two single-shank Neuropixels^60^ probes (5 rats) in the left mEC and right CA1 region (the surgical procedures described above). During a trial, each rat would forage in one of the two enclosures for at least 60 minutes; each experimental session consisted of two trials in different enclosures. The rat’s position was determined based on an array of two infrared LEDs fixed to the rat’s head as described above (sampled at 50 Hz). Speed was smoothed using a moving average (box) filter with a span of 20 samples (400 ms).

It should be noted that the large-arena datasets were included only in the analysis relating to field scale and time shifts to include grid and place cells with as large scale/field sizes as possible (Fig. 4). These large-arena recordings did not include simultaneously recorded grid cells of multiple scales or the hippocampal LFP; hence it this new data could not be used for the analysis in Figs. 5-6. Furthermore, we restricted our results to the data from^11^ in Figures 1 and 2 to avoid substantial differences in field sizes between the grid and place-cell populations and to enable direct comparison between the prospective temporal properties and the results on spatial deformations reported in^11^.

### Rate maps and smoothing

We divided each enclosure area into 2.6 × 2.6 cm bins and then used the recordings to make maps of (a) the number of spikes fired in each bin and (b) the time spent in each bin (“dwell time”). We smoothed each of these maps by convolution with a Gaussian kernel and produced each smoothed rate map by dividing binned spike counts by dwell times. We defined the kernel width as σ = l / 2π, where l was the position of the first trough in the radial average SAC. Place cells were smoothed with a fixed kernel width of σ = 3 bins (7.9 cm). Smoothed rate maps were used for all subsequent methods unless specified otherwise. Unvisited bins within the bounds of the map are shown in white in Fig. 1A. For all ZLAC calculations, these bins were assigned rates by interpolating between adjacent bins.

### Cell classification

Grid cells were identified as cells with a gridness score of > 0.27^11,61^. Only non-directional grid cells were used in the current study to make sure that the results are not due to directional bias. Only CA1 cells with clearly defined fields were classified as place cells and included in this analysis as described in^11^.

We identified conjunctive grid cells^62^ and similar direction-sensitive place cells by calculating the head direction (HD) score of each recording^63^. We produced a polar histogram showing each cell’s firing rate as a function of HD, using 1° bin and smoothing the resulting counts with a rolling 25-bin box filter. To assess the directionality of this distribution, we calculated the Rayleigh vector as follows:

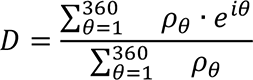

where ρ_θ_ represents the polar firing rate as a function of HD in the bin [θ-1, θ].

This vector’s magnitude, |D|∈[0,1] was used as the HD score. Cells with |D| > 0.5 during any of the trials were excluded from subsequent analysis. This procedure and the thresholds used are identical to those used by^63^.

### Spatial autocorrelation (SAC) and cross-correlation (SCC)

Let *λ*_1_ and *λ*_2_ denote two unsmoothed firing rate maps, where *λ* (*x*, *y*) is the value of rate map *λ* at the coordinates (*x*, *y*). The normalised spatial cross-correlation (SCC) between these maps is calculated for each possible discrete spatial lag (*τ_x_*, *τ_y_*) (measured in bins) as follows, as defined by^25^:

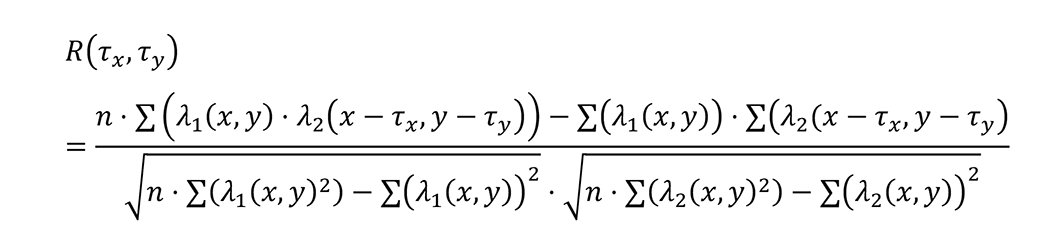

where each summation *Σ* is over all *n* bins for which rates were estimated in both *λ*_1_ and *λ*_2_. Setting *λ*_1_ equal to *λ*_2_ makes the above an equation for spatial autocorrelation (SAC). Spatial cross-correlograms were smoothed with a Gaussian kernel (*σ* = 2 bins, or 5.2 cm), before subsequent analysis.

### Zero-lag autocorrelation (ZLAC)

The formula defined above is equivalent to calculating Pearson’s correlation coefficient of a rate map with a copy of itself at different spatial lags; hence, *R* (0, 0) = 1. To measure the ‘sharpness’ of each rate map, we used the following simplified version of the autocorrelation equation:

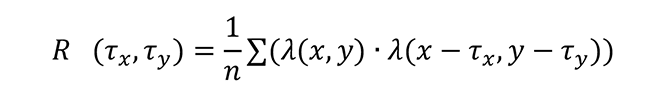

This version of autocorrelation removes the normalisation usually applied to a spatial autocorrelogram. This means that the autocorrelation value at zero spatial lag, i.e. R_sim_ (0, 0), is no longer guaranteed to be equal to 1. Instead, when the time-shifted version of a rate map has a higher R_sim_ (0, 0) than the original, it can be inferred that the map’s fields are sharper than before. This is equivalent to the mean squared value of each rate map. Each map’s optimum time-shift was determined as the position of the ZLAC peak nearest 0 ms. These correlograms were smoothed with the same Gaussian kernel used for normal SACs (σ = 2 bins, or 5.2 cm).

### Radial average SAC

We determined each grid cell rate map’s scale by calculating its radial average SAC. These were calculated by averaging of all bins within shells of increasing radius l from the smoothed autocorrelogram:

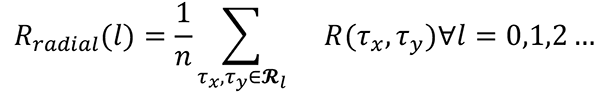

where each summation is over the set 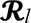 of all *n* bins where 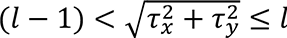. To find the grid cell’s scale, *R*_radial_ (*l*) was smoothed using a moving average (box) filter over a span of 9 bins; the grid cell’s scale was taken as the position of the first peak at *l* > 0.

### Spatial Information

We calculated the spatial information *S* using a method similar to^20^:

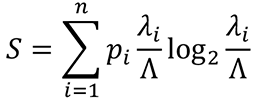

where Λ is the mean firing rate of the cell; *λ*_*i*_ is the mean value of bin *i* in the smoothed firing rate map; and 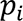 is the mean value of bin *i* in the smoothed map of dwell times. Each map is divided into spatial bins *i* = 1, … *n*. Note that we use bin values from the smoothed versions of each map.

### Time shifting

We used the following procedure to “time shift” recording data. During each trial, the rat’s position and head direction were recorded at 50 Hz. Hence, we organised the spikes fired by each cell into a vector of spike counts, also sampled at 50 Hz. This spike count vector was shifted by a given number of samples to lead or lag the position and head direction vectors (leading = “positive”, lagging = “negative” time shifts). Each sample corresponds to a shift of 20 ms; recordings were time-shifted by up to ±2 seconds. Rate maps were then recalculated from these vectors as normal, using the original smoothing parameters.

### Validation of ZLAC analysis of time shifts

We observed a significant correlation between cells’ optimal time shifts and the size of their firing fields. To identify any methodological bias, we generated grid-and place-cell activity patterns with scales/field sizes matching the real data (Fig. S9A-B; see sections below). We combined these with randomly selected trajectories from the actual dataset to simulate several spike counts and the resulting rate maps (Fig. S9C-D; see sections below). We then used the same ZLAC analysis applied to the experimental data to test whether identified ZLAC peaks showed any consistent deviation from the correct time shift of 0 (Fig. S9E). We found that the median identified time shift was more variable with increasing field size (Fig. S9F-G). However, there was no evidence to suggest that the ZLAC analysis showed a bias towards non-zero median shifts when firing fields were larger (Benjamini-Hochberg test, α = 0.05).

### Choosing place cells with single fields

When analysing the relationship between place-field size and time shifts, we restricted our analysis to cells with single fields to simplify interpretation. We found the rates (λ) and positions of all local maxima on the place cell’s rate map after smoothing, including the bin with the highest overall rate (λ_max_). If any of the other local maxima had rates λ>0.4·λ_max_, we classified that map as containing multiple potential fields and excluded it from this particular analysis. Each field area was defined as the area of pixels with rates λ>0.2·λ_max_ surrounding the highest-rate bin.

### Simulating grid cell activity functions

We modelled grid-cell firing fields as three overlapping 2D plane waves oriented at 60° offsets to each other (Fig. S9A). First, we calculate the total wave vector, k:

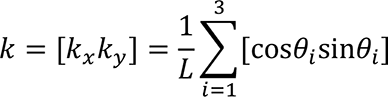

Each wave has wavelength 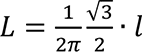, where *l* is the grid scale. For a given point (*x*, *y*), the grid cell’s activity *f*_*x*,*y*_ is calculated as follows:

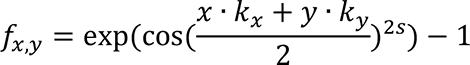

The parameter s determines the ‘sharpness’ of the resulting grid; higher values decrease the width of each firing field. For all simulated grid cell rate maps, s=0.5.

### Simulating place cell activity functions

We modelled place cell firing fields as 2D Gaussian functions (Fig. S9B), i.e. for a given point x=[x,y], the place cell’s activity is calculated as

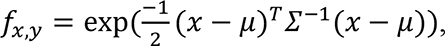

where the vector μ represents the coordinates of the place field centre, and *Σ* = [*σ*00*σ*]. The parameter *σ* controls the width of the resulting place field, which is circular.

### Drawing spike vectors from simulated rate maps

The activity functions described above specify the probability that a grid cell will spike in a particular position (as the rate parameter of an inhomogeneous Poisson distribution). To simulate a grid cell’s firing, we specified its activity function, selected a trajectory taken by a rat during a randomly selected trial, and used the function to assign a probability of spiking to each point along this trajectory. We then assigned a fixed number of spikes to points along this trajectory in line with said probabilities (grid cells: 2000 spikes, place cells: 500 spikes) (Fig. S9C-D).

We simulated grid cell rate maps that varied in their scale (0.3 to 1.2 m, in 0.3 m steps) and place cell rate maps that varied in the size of the (single) firing field (covering 0.1 to 0.4 m^2^, in 0.1m^2^ steps). Grid cell rate maps varied in their phase offset, so all possible horizontal and vertical phase offsets of each specified map were uniformly sampled. Similarly, we randomised the position of the field on each place cell rate map. Each combination of scale and time shift was sampled 2000 times.

### Defining individual firing fields and runs through fields

To study time shifts on single runs through grid cells’ firing fields, we segmented each rate map into fields by applying MATLAB’s inbuilt watershed function to the negative rate-map. This identifies each minimum’s “drainage basin”; each identified basin corresponds to a field surrounding a peak in the original rate map. To avoid over-identifying spurious peaks as potential field centres, we applied MATLAB’s extended minima transform to remove minima whose depth was less than σ/5, where σ is the standard deviation of all of the image’s pixels. We excluded fields whose peaks were less than 15 cm from the edge of the enclosure. We removed pixels on the edges of each field whose firing rates were <20% of its peak firing rate. This 20% threshold was also used for defining each place cell’s total firing field area.

The rat’s trajectory was divided into segments, or “runs”, passing through each field. We assigned each spike on this run (Fig. 2D, red dots) an individual, ‘single-run’ time shift relative to the point on the run passing closest to the field centre. Spikes fired before (after) this point were assigned positive (negative) time shifts. The overall time shift for the run was defined as the median of these time shifts (Fig. 2D, green dots). We only considered runs that (a) had an associated spike train and (b) passed within half the field radius of the field’s centre.

### Investigating time shifts of spike trains when approaching or leaving walls

We divided each enclosure into inner and outer zones of equal area (Fig. 2F). We focused on fields where ≥30% of each field fell within the outer zone (called peripheral fields). We measured the distance between each bin on the edge of the peripheral field and the closest bin on the enclosure boundary and used these distances to divide these bins into “inner” and “outer” groups of equal size, where the “outer” group represents the half of the field’s circumference closest to the wall (Fig. 2F-G). We then isolated valid runs (see criteria above) that started in the inner group and ended in the outer group or vice versa and labelled these runs “outward” or “inward” runs, respectively. For each map, we collated the time shifts of inward and outward runs across all valid peripheral fields. We excluded grid-cell maps with fewer than 15 inward or 15 outward runs in total. As place cell fields covered a smaller proportion of each arena, we reduced the minimum numbers of inward and outward runs to 5 each.

In each recording with sufficient runs, we looked at the median time shift on inward and outward runs and the difference between them. The final value assigned to each cell was the grand median of these values for all trials in which said cell was recorded during a given session.

### Comparing time shifts of spike trains from pairs of co-recorded grid cells

To estimate the difference between time shifts in overlapping grid fields, we calculated all the median time shifts of all individual laps through the fields. The runs from different cell pairs were considered overlapping if they occurred within 200 ms from each other and the distance between the average respective field centres was less than 5 bins (i.e. < 13 cm). We then calculated the time-shift differences between all co-recorded runs of all co-recorded grid-cell pairs. We split these differences in time shifts (larger field - smaller field) into intra-module (grid scale ratio <1.1) and inter-module (grid scale ratio ≥ 1.4), which were compared using a two-sample t-test. We also used Student’s t-test to test whether the time shift differences of inner-grid modules differed significantly from zero.

### Analysis of external and internal cues’ effects

When regressing time-shift (Δ*t*) against enclosure shape (Fig. 2A) and heading (Fig. 2B), we removed per-cell (and per-subject) effects by standardising time-shifts across cells. We replaced the per-cell median Δ*t* with the population median Δ*t* before aggregating. We regressed time-shift (Δ*t*) against direction *θ* by fitting Δ*t* = *a* cos(*θ*) + *b* sin(*θ*) + c using least-squares. This is equivalent to the model Δt = *d* cos(*θ* – *θ*_max_) + c, where the modulation depth “*d*” is given by *d*² = *a*² + *b*², and *θ*_max_ = tan^-1^(*b* / *a*) is the direction in which time shifts tended to be the largest. We assessed significant tuning by re-fitting the model on shuffled *θ*, and calculated confidence distributions for *d* and (differences therein) by bootstrap resampling (1000 samples).

We calculated per-run acceleration (angular speed) as the change in forward speed (head direction) during a run through a single field divided by the run’s duration. We calculated per-run heading (head directions, forward speeds) as the average heading (head direction, forward speed) throughout every single run. We calculated Spearman’s rank correlations between time shifts and per-run acceleration, forward speed, and angular speed. To remove per-cell effects, we converted time shifts to ranks before aggregating. To remove per-subject effects, we converted acceleration and speed data to ranks per-subject before aggregating. To invert the rank transform when plotting Fig. 3A, we took the average time-shift (over aggregated cells) corresponding to each rank (in bin sizes of two percentage points), and smoothed the resulting function with a Gaussian (σ: 2% percent) to remove further variance. To calculate partial correlations, we used the Python package 'Pingouin'^64,65^

### Theta analysis

The LFP signal was recorded in the hippocampus using one of the tetrode wires. It was amplified 2000–8000 times, bandpass filtered at 0.34–125 Hz and sampled at 250 Hz. The LFP signal was only available for analysis from multi-tetrode recordings. The signal phase was calculated as the argument of the Hilbert transform H (t) (MATLAB).

### Pairwise phase consistency

We restricted our analyses of theta phase-precession to cells that showed significant coupling to the theta rhythm, as assessed by phase-locking statistics. The theta phase-locking value for a given recording was calculated as the magnitude of the average complex-valued phase-vector (“phasor”):

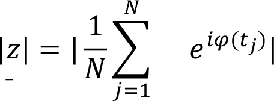

where *t*_*j*_ is the time at which spike *j* was fired, *N* is the number of spikes fired by the cell during the trial, and *φ*(*t*) is the theta phase in radians. To decrease the bias induced by finite sampling, we calculated the pairwise phase consistency (PPC, or *Ŷ*; ^66,67^), which corrects for this bias:

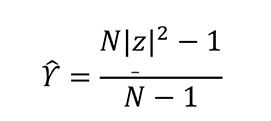

For large 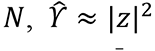. We assessed each recording’s degree of theta phase locking by comparing *Ŷ* to a shuffled distribution, produced by circularly shifting the recorded LFP signal 2000 times before recalculating *Ŷ* (minimum offset 10 seconds in either direction). Any recordings with a *Ŷ* value that exceeded the 99^th^ percentile of their respective shuffled distributions were deemed to display significant phase locking, and subsequent in analyses correlating preferred theta phase with time shifts.

### Assessment of correlations

Unless otherwise stated, we evaluated correlations using Spearman’s rank correlation coefficient (ρ), which is appropriate when the underlying correlation is not necessarily linear. There were three major exceptions to this. We used Pearson’s correlation coefficient to evaluate the correlation between optimum shifts and the inverse of the running speed and the correlation between single-run time shifts and ‘fixed’ time shifts estimated by ZLAC (Fig. 2C), since any correlations were expected to be linear in both cases. We calculated correlations between preferred theta phases and time shifts using Kempter’s circular-linear correlation coefficient (CLCC)^68^, since the phase is a circular variable.

## Data availability

The full raw dataset is available from the corresponding author on reasonable request.

## Code availability

All custom code written for reported analyses are publicly available at [insert Github link] or via request to the corresponding author.

## Acknowledgements

P.C.V. is supported by MRC, Frank Elmore Fund, and University of Cambridge School of Clinical Medicine. M.R. is supported by a Leverhulme Trust Fellowship and HFSP grant ECF-2020-352. M.B. is supported by the Wellcome Trust Grant (100154/Z/12/A). M.K. is supported by BBSRC DTP at University of Cambridge. T.O’L. is supported by ERC grant 716643 FLEXNEURO and HFSP grant RGY0069/2017. J.K. is a Wellcome Trust/Royal Society Sir Henry Dale Fellow (206682/Z/17/Z) and is supported by Dementia Research Institute (DRICAMKRUPIC18/19), Isaac Newton Trust/Wellcome Trust ISSF/University of Cambridge Joint Research Grant, Kavli Foundation Dream Team project (RG93383), Isaac Newton Trust [17.37 (t)], and NVIDIA Corporation.

The funder had no role in study design, data collection and analysis, decision to publish or preparation of the manuscript. For the purpose of open access, we have applied a CC BY public copyright license to any author-accepted manuscript version arising from this submission.

## Author contributions

J.K., M.B. and T.O’L. conceived the study. S.B., M.B., M.K., P.K. and J.K. collected the data. P.C.V. and M.R. analysed the data with contributions from M.B., T.O’L. and J.K. J.K, M.R. and P.C.V. wrote the manuscript with contributions from M.B. and T.O’L.

## Corresponding author

Correspondence and requests for materials should be addressed to J.K.

