## Supplementary Information for "Simultaneous representation of multiple time horizons by entorhinal grid cells and CA1 place cells"

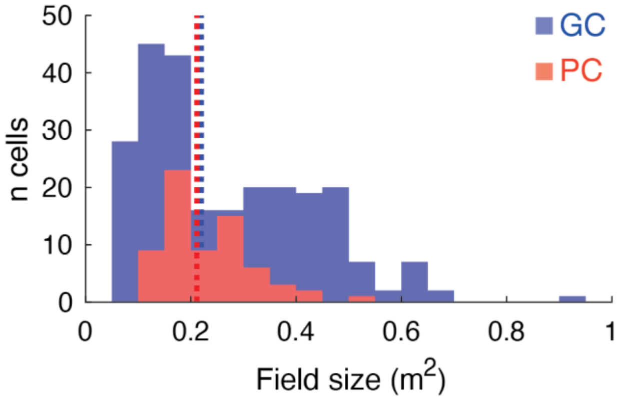


**Supplementary Figure 1 (related to Figure 1). There is no significant difference between the median field sizes of grid cells (blue) and place cells (red)**. The median field size for each cell type is indicated by dotted lines of matching colours.

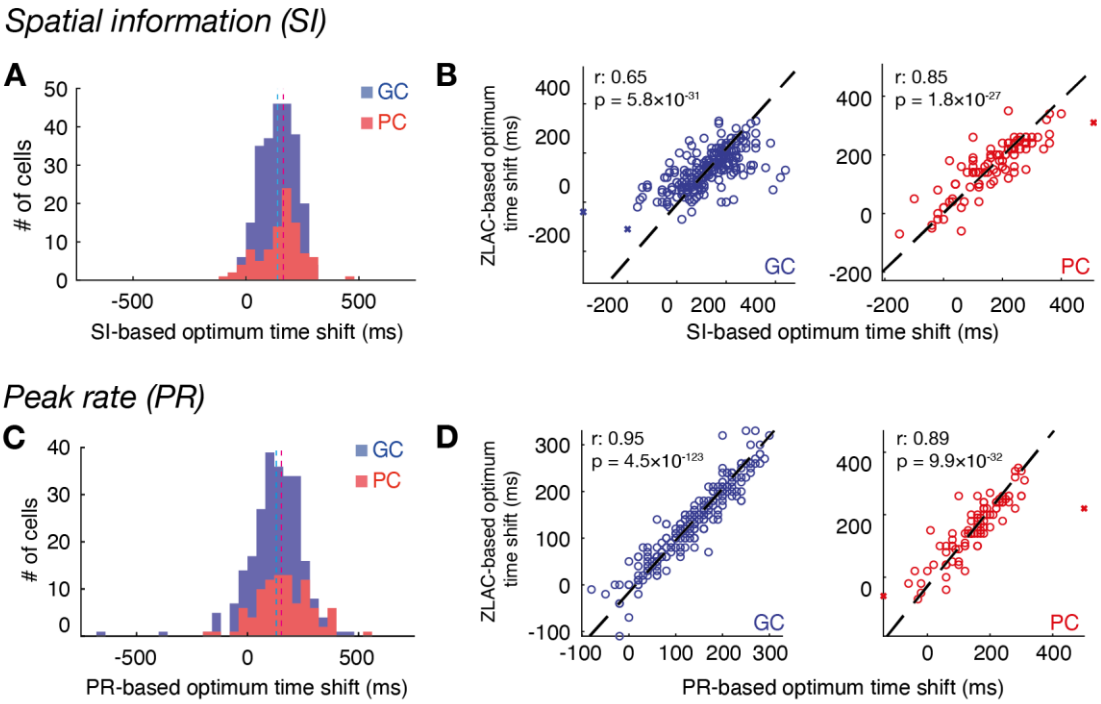


**Supplementary Figure 2 (related to Figures 1 and 2). Zero-lag autocorrelation (ZLAC) as a measure of rate map sharpness is comparable to other measures used for place cells.** To validate ZLAC as a measure of rate map sharpness, we re-calculated each cell's optimum time shift (median across each trial in which a cell was recorded) by optimising each rate map's spatial information (SI, upper row) and peak rate (PR, lower row). Histograms (**A**, **C**) show distributions of optimum time shifts as calculated with SI (**A**) and PR (**C**). Scatter plots (**B, D**) compare each cell's time shift as calculated by ZLAC (as in Fig. 1C; y-axis) and by SI (**B**, x-axis) or PR (**D**, x-axis). GC: grid cell; PC: place cell.

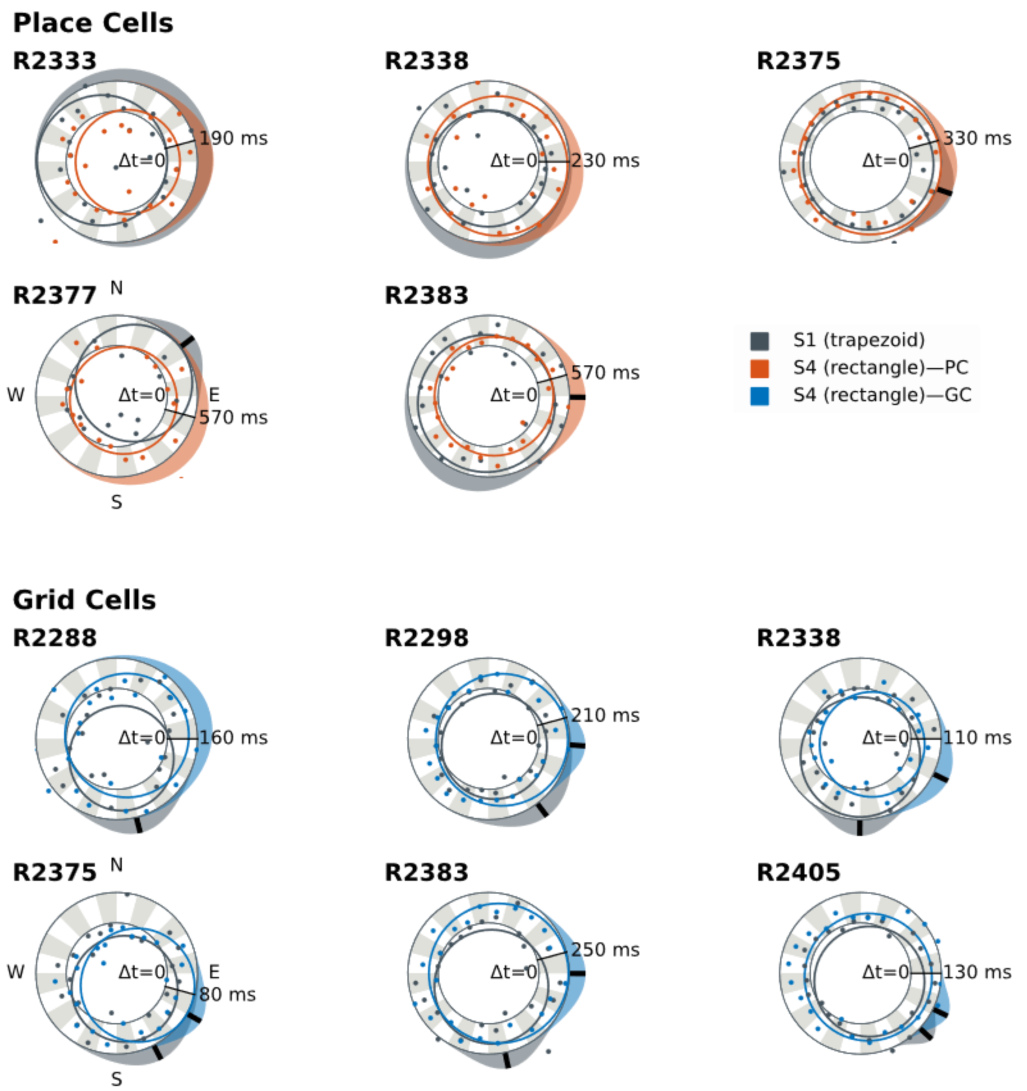


**Supplementary Figure 3 (related to Figure 2). The heading direction in which time shifts rotate in response to the deformation of arena walls.** The directional dependence of per-run time shifts (Δt) on heading direction for place (top, red) and grid cells (bottom, blue). Each log-polar plot shows a heading on the angular axis and a time shift on the radial axis (inner ring: $\Delta t = 0$). Plotted lines show the best-fit polar least-squares regression $\Delta t = a \cos(\theta) + b \sin(\theta) + c$, relating heading direction θ to Δt; Points show the median time shift in 15° bins (lines: 95% confidence) for the trapezoidal arena ("S1"; black) and rectangular arena ("S4"; coloured). Shaded distributions on the exterior show von Mises confidence intervals for the direction in which shifts are largest ($\theta_{max}$). Black bars for $\theta_{max}$ are plotted fits that remained significant after two-stage Benjamini-Hochberg correction for a false-discovery rate α=0.05. Place-cell sampling was too sparse to draw firm conclusions. In grid cells, the heading of maximal Δt rotated to reflect the orientation of the west wall of the enclosure (c.f. Fig. 1A).

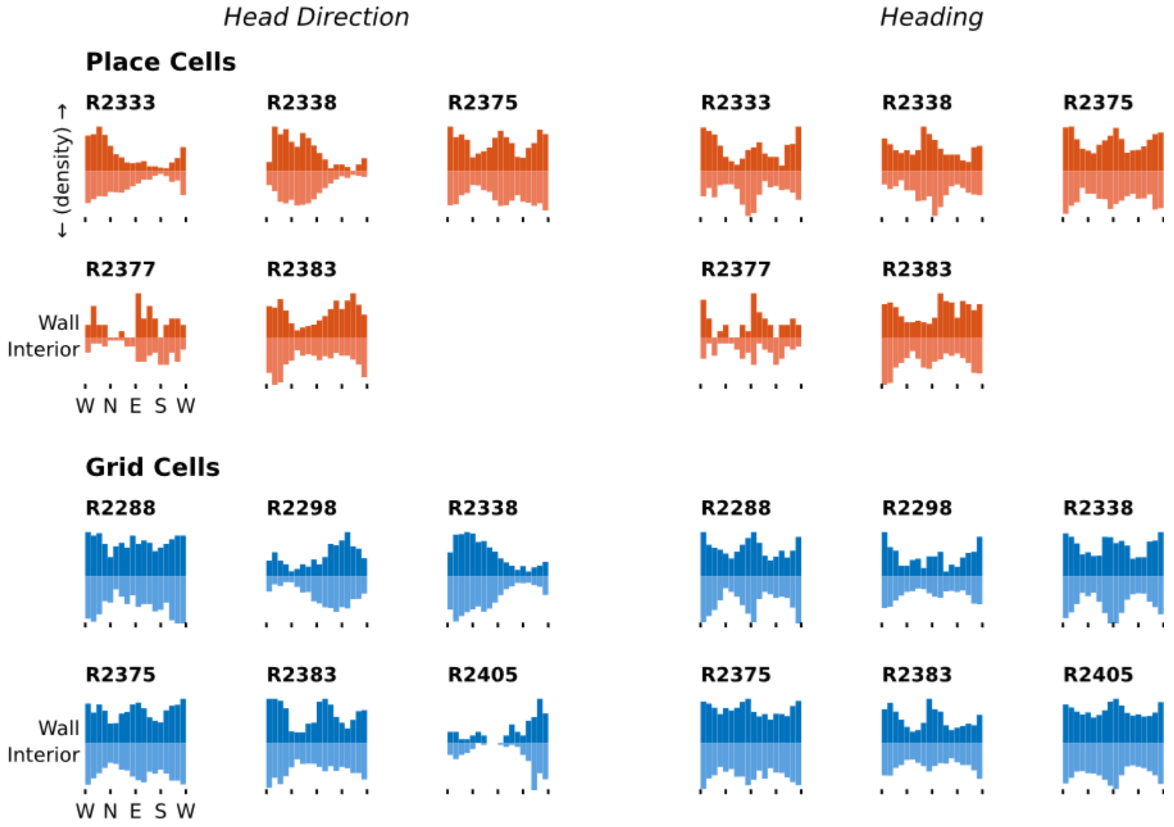


**Supplementary Figure 4 (related to Figure 2). Heading direction coverage: interior vs exterior.** Each plot shows the density of single-run head-direction angles (left) and heading angles (right) for a single rat, for fields near walls (above) and in the interior (below).East–west travel directions are over-sampled, as expected for the same of the arena, but there are no overt differences between the travel directions in the exterior or interior. Note that head direction and heading show similar distributions, therefore, we used only the latter in the analysis describing the influence of internal cues.

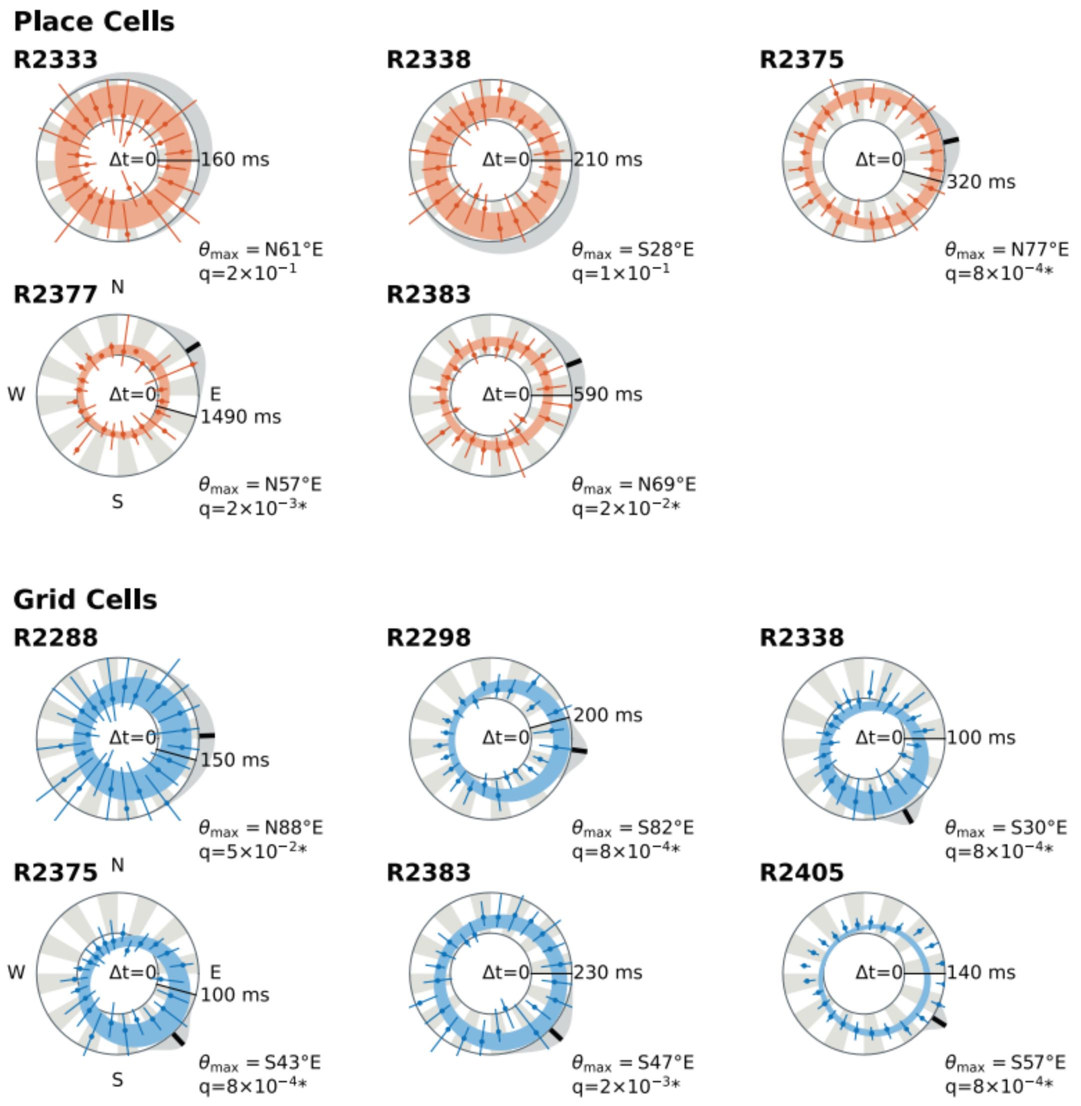


**Supplementary Figure 5 (related to Figure 2). Time shifts are more prospective when the animal is heading eastwards.** The directional dependence of per-run time shifts (Δt) on heading direction for place (top, red) and grid cells (bottom, blue). Each log-polar plot shows a heading on the angular axis and a time shift on the radial axis (inner ring: $\Delta t = 0$). Shaded regions show the 95% confidence interval for a polar least-squares regression $\Delta t = a\cos(\vartheta) + b\sin(\vartheta) + c$ relating heading direction $\vartheta$ to $\Delta t$; points show the median time shift in 15° bins (lines: 95% confidence). Heading modulation was significant (*) after false-discovery-rate correction (α=0.05, two-stage Benjamini-Hochberg) in place cells in three subjects and grid cells in all subjects. Distributions plotted on the exterior of the ring show a von Mises confidence interval for the direction of maximal Δt, $\vartheta_{max}$.

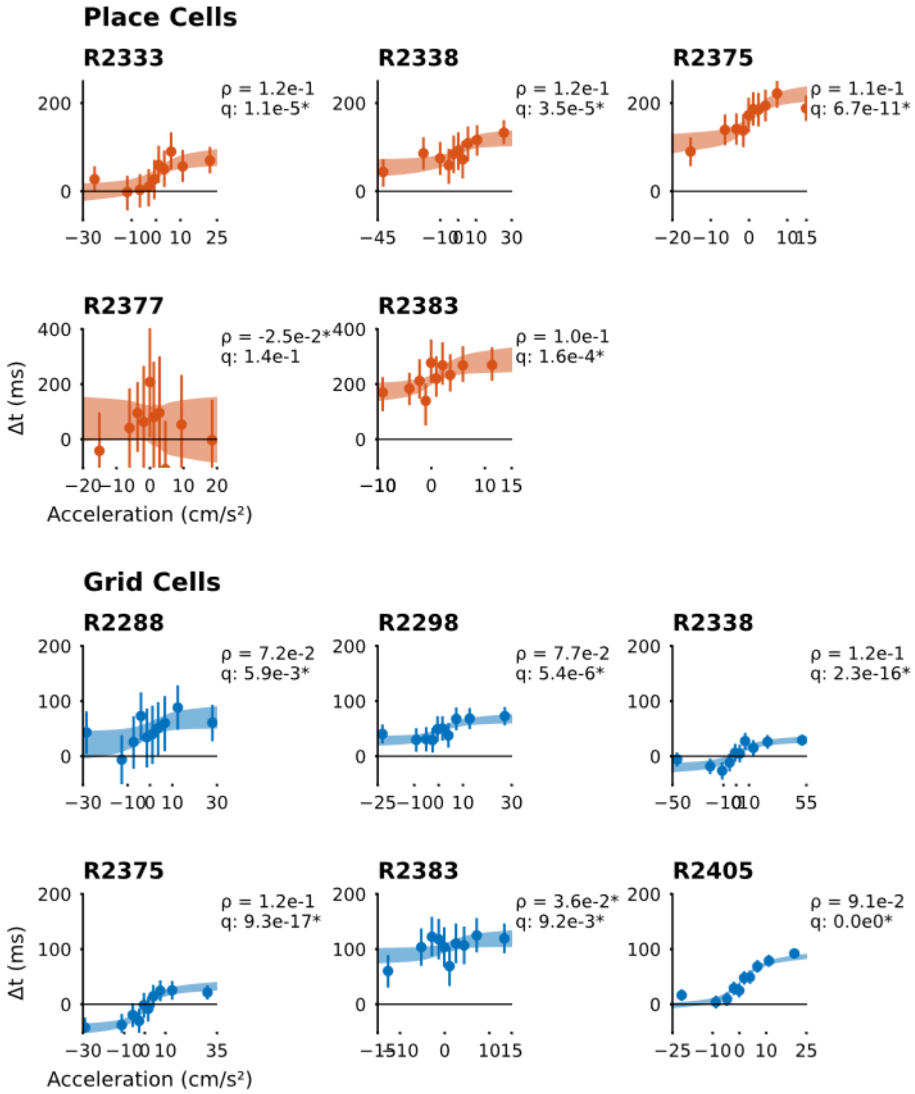


**Supplementary Figure 6 (related to Figure 3). Time shifts weakly but positively correlated with acceleration.** We used a rank regression to correlate single-run time shifts with the average acceleration for each run. Plots show the regressed model transformed into physical units for illustration (red: place cells, blue: grid cells). The shaded regions reflect the 2.5–97.5th percentile confidence interval for the linear fit after transformation. Points show the mean time shifts for each decile of acceleration (lines: bootstrapped 95% confidence intervals for the mean time shift within each decile). The residual variance in Δt is so large that we cannot visualise it cleanly on these plots. We converted per-run acceleration to ranks for each rat (all sessions combined), and per-run times shifts to ranks for each cell separately. We combined all cells from each subject and fit a linear least-squares regression to predict ranked Δt from ranked acceleration. The fraction of explained variance $R^2$ for this regression is the square of Spearman's correlation coefficient. With the exception of place cells for R2377, all correlations were significant after correcting for a 5% false discovery rate via two-stage Benjamini-Hochberg. Nevertheless, acceleration explained only a fraction of the variance in time shifts. ρ: Spearman correlation coefficient; q: two-stage Benjamini Hochberg false-discovery corrected p-value.

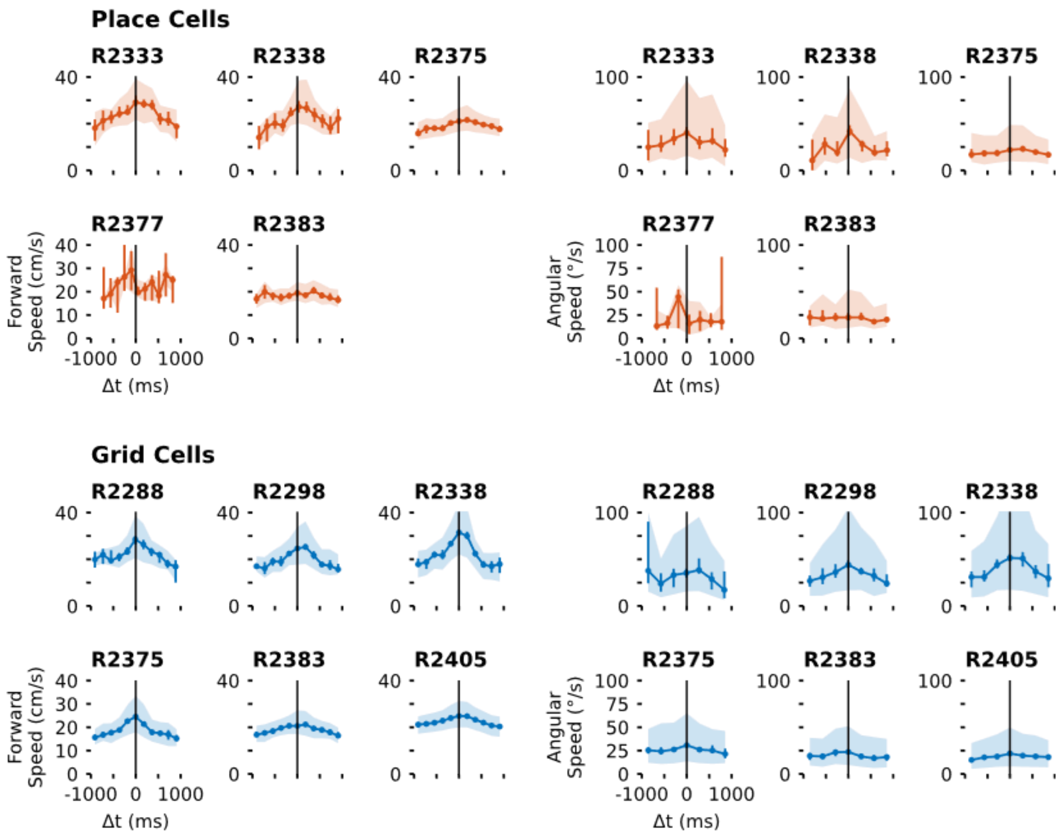


**Supplementary Figure 7 (related to Figure 3). Speed shows a nonlinear relationship to time shifts that is consistent across subjects.** Both forward (running) speed **(left)** and the angular speed of head direction **(right)** tend to be smaller when time shits (Δt) are extremely positive or negative. Larger speeds are associated with smaller time shifts. Plots for each subject and cell type show the relationship between the speed and time shifts on single runs through a given place **(top)** or grid **(bottom)** field. Points show the median speed within a histogram bin for Δt (error bars: 95% bootstrapped confidence intervals). Shaded regions show the interquartile range (the full variation is too large, relative to the median trend, to visualise in a single plot).

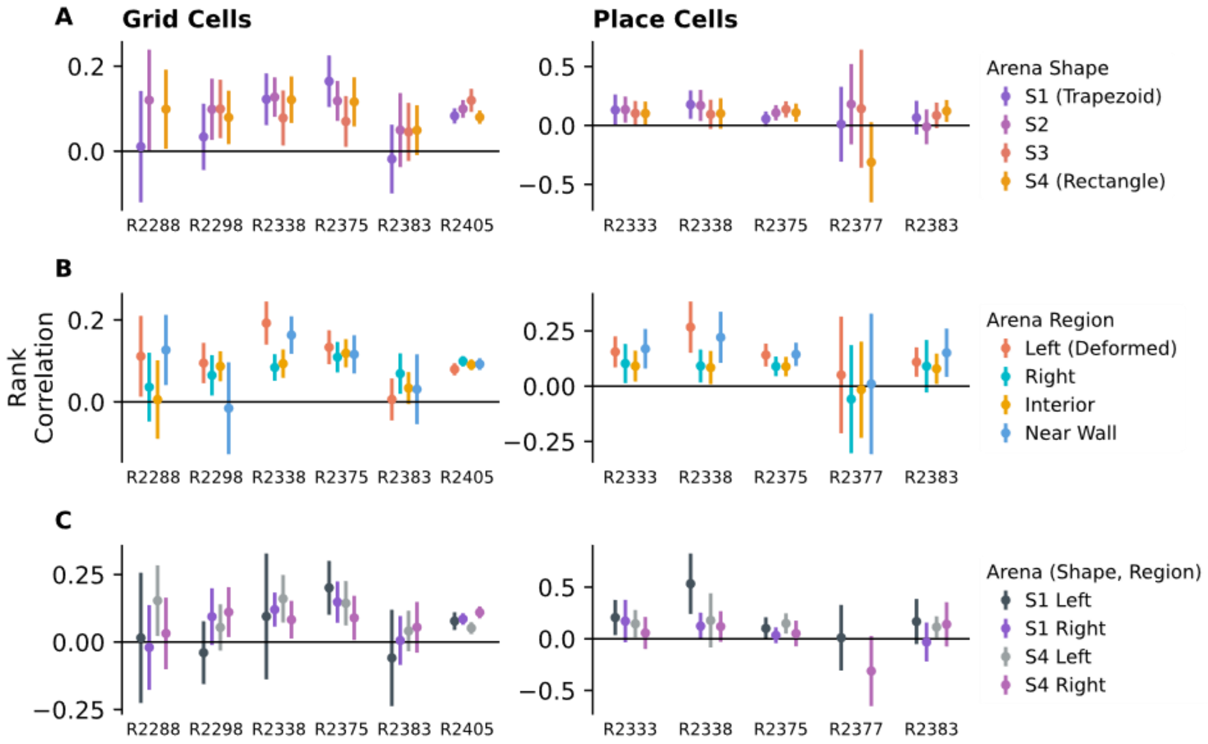


**Supplementary Figure 8 (related to Figure 3). The dependence of time shifts on per-run acceleration does not show strong spatial topography. (a)** Nonlinear (Spearman) correlation between acceleration and time shift broken down by the shape of the arena (lines: 2.5–97.5% confidence). No consistent trends are visible. **(b)** The same, broken down by region of the arena. Again, no consistent trends emerge. **(c)** Comparing the left half of the trapezoidal (S1) arena to the non-deformed right half of S1 or either half of the rectangular S4 showed no consistent trends. We conclude that acceleration modulates prospective time shift weakly and independently of, e.g. encountering walls on the periphery. Missing bars indicate conditions with too few single-trial runs to compute correlations.

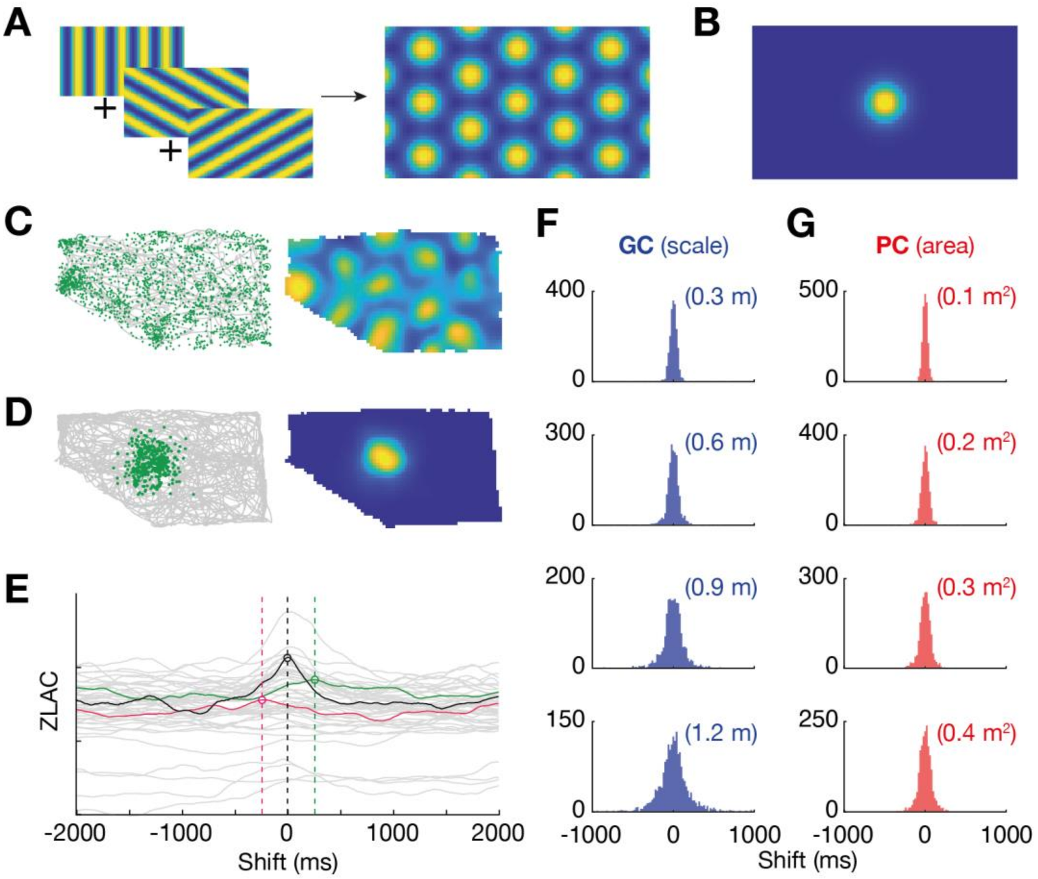


**Supplementary Figure 9 (related to Figure 4). In the simulated grid and place cell rate maps, there is no correlation between firing field size and the time shifts identified by ZLAC analysis. A** Grid cell activity was modelled by taking the sum of three plane waves oriented at 60° offsets to each other and then adjusting the sharpness of the resulting grid. **B** Place cell activity was modelled as a 2D circularly symmetric Gaussian function. **C** Example of spikes drawn from the grid cell activity map in (a) and assigned to various points along the trajectory (left) and the resulting rate map (right). **D** Example of spikes drawn from the place cell activity map in (b) and assigned to various points along the trajectory (left) and the resulting rate map (right). **E** ZLAC of 50 simulated grid cell spike vectors with respect to time shifts (grey lines). Each is drawn using the activity map shown in (a) (right); this map is used to randomly assign 2000 spikes to points along a rat's trajectory randomly selected from the real data. Here, though the average optimum time shift is zero (e.g. black line), some spike vectors drawn with this method may show negative (magenta) and positive (green) optimum time shifts. **F** Histogram showing the optimum shifts of 2000 simulated grid cell spike vectors. None of these medians showed any significant deviation from 0 ms after correction for multiple comparisons (Benjamini-Hochberg test, α = 0.05). **G** Histogram showing the optimum shifts of 2000 simulated place cell spike vectors. None of these medians showed any significant deviation from 0 ms after correction for multiple comparisons (Benjamini-Hochberg test, α = 0.05). GC: grid cell; PC: place cell.

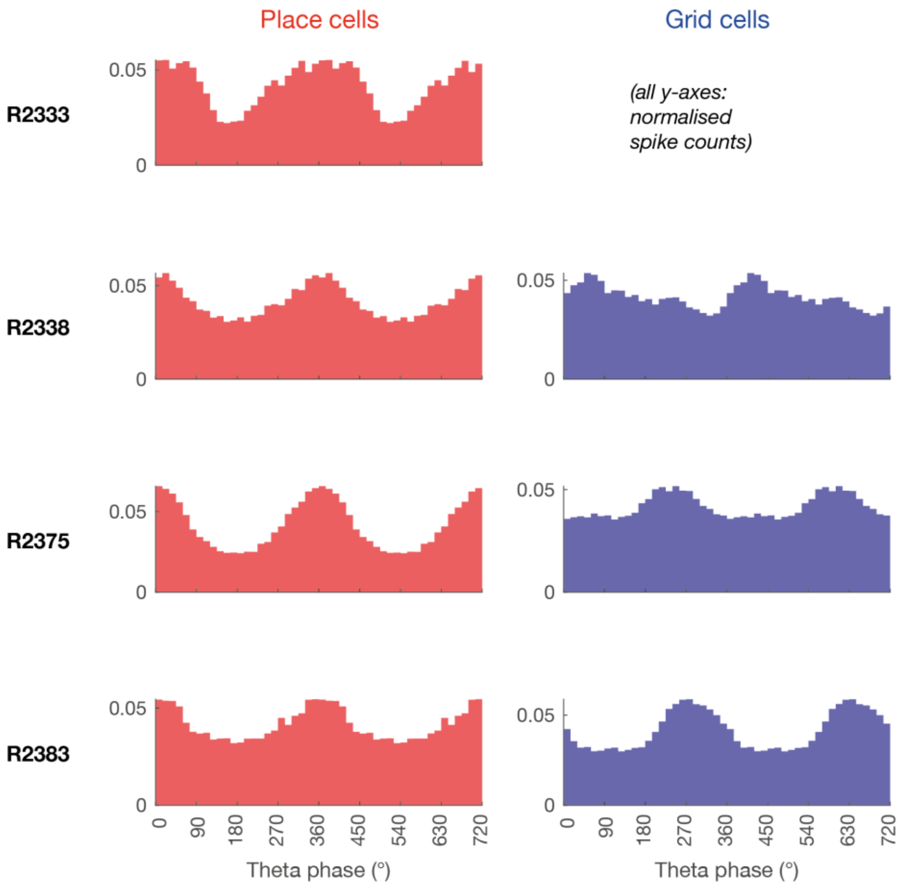


**Supplementary Figure 10 (related to Figure 6). Histograms showing normalised spike counts with respect to *stratum pyramidale* theta phase,** grouped by the animals in which they were recorded (place cells: red, left; grid cells: blue; right). Note that no grid cells were successfully recorded from R2333.

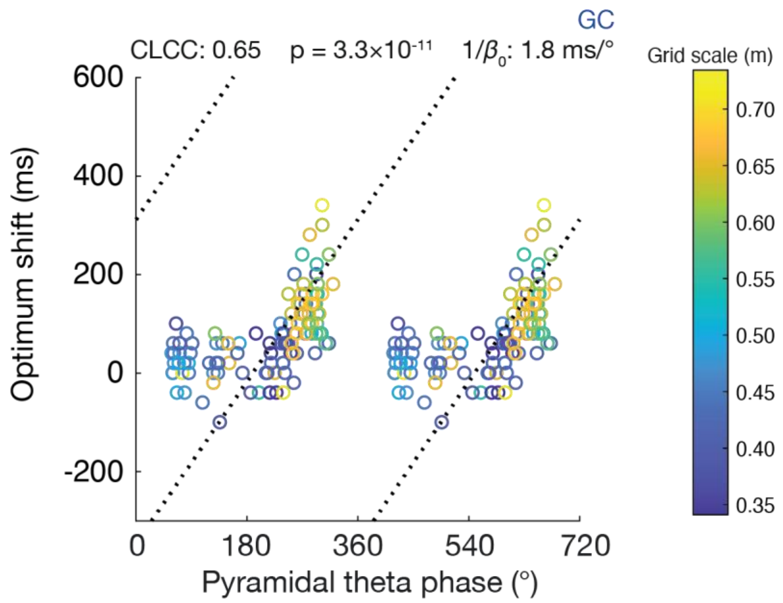


**Supplementary Figure 11 (related to Figure 6). Larger-scale grid cells tend to have larger optimum time shifts.** Scatter plot showing each grid cell recording's preferred pyramidal theta phase against its optimum time shift; the colour map shows the grid scale measured in each map. Note that cells with large scales (~70 cm; yellow) concentrate at larger shifts and later theta phases, compared to cells with small scales (~35 cm; blue). GC: grid cell; CLCC: circular-linear correlation coefficient; $\beta_0$: slope parameter of the regression line (units ° phase/ms).

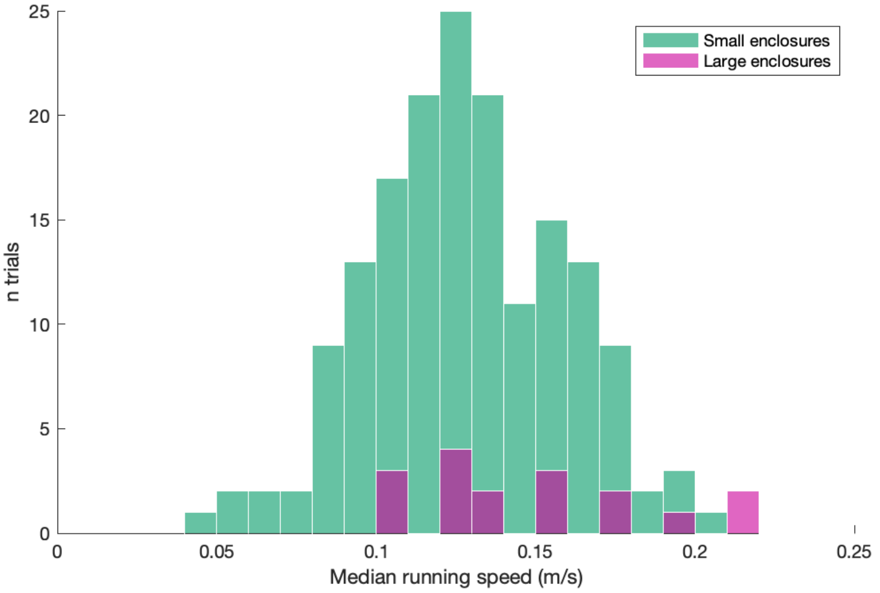


**Supplementary Figure 12 (related to Figures 4 and 5).** Median running speeds across all trials in which cells were recorded in smaller enclosures (“S1”-“S4”, green) and larger enclosures (magenta) (all subjects combined).

**Table S1. Comparison of time shifts in cells with/without significant theta precession.** Missing values indicate rats with no cells of the specified type. Blank rows indicate animals for which LFP was not recorded from the hippocampus. Overall, we could not detect a significant difference between the time shifts of precessing and non-precessing cells.

| | PLACE CELLS | | | | GRID CELLS | | | |
|---|---|---|---|---|---|---|---|---|
| | Precessing | | Non-precessing | | Precessing | | Non-precessing | |
| | No. of cells | Median optimum shift (ms) | No. of cells | Median optimum shift (ms) | No. of cells | Median optimum shift (ms) | No. of cells | Median optimum shift (ms) |
| R2288 | -- | **--** | -- | **--** | 3 | **60** | 1 | **120** |
| R2298 | -- | **--** | -- | **--** | -- | **--** | -- | **--** |
| R2333 | 5 | **130** | 2 | **83** | -- | **--** | -- | **--** |
| R2338 | 3 | **160** | 5 | **190** | 4 | **52** | 5 | **30** |
| R2375 | 18 | **180** | 14 | **200** | 10 | **40** | 3 | **10** |
| R2377 | -- | **--** | -- | **--** | -- | **--** | -- | **--** |
| R2383 | 2 | **153** | 7 | **213** | 12 | **118** | 1 | **220** |
| R2405 | -- | **--** | -- | **--** | -- | **--** | -- | **--** |
| All | 28 | **165** | 28 | **192** | 29 | **67** | 10 | **33** |
| Precessing vs. non-precessing | | | | **(p = 0.23)** | | | | **(p = 0.31)** |

**Table S2. The circular-linear correlation coefficient between recordings' preferred pyramidal theta phase and their optimum time shifts, split between animals.**

| | PLACE CELLS | | GRID CELLS | |
|---|---|---|---|---|
| Rat ID | No. of recordings | **CLCC** | No. of recordings | **CLCC** |
| R2288 | -- | **--** | -- | -- |
| R2298 | -- | **--** | -- | -- |
| R2333 | 32 | **0.37** | -- | -- |
| R2338 | 31 | **0.14** | 30 | **0.29** |
| R2375 | 144 | **0.31** | 45 | **0.64** |
| R2377 | -- | **--** | -- | -- |
| R2383 | 32 | **0.67** | 42 | **0.35** |
| R2405 | -- | **--** | -- | -- |
| All | 239 | **0.36 ($p = 3.6\times10^{-6}$)** | 117 | **0.65 ($p = 3.3\times10^{-11}$)** |

Recordings were only included if they showed significant theta phase locking (as measured by PPC – see Methods) when compared to a shuffled distribution. Note that this table only includes data from animals with both (a) CA1 place cell recordings and (b) LFP recorded in the hippocampus.
